# Blue Light Sonata: Dynamic variation of red:blue ratio during the photoperiod differentially affects leaf photosynthesis, pigments, and growth in lettuce

**DOI:** 10.1101/2025.01.24.634700

**Authors:** Jordan B Van Brenk, Kimberly R Vanderwolk, Sumin Seo, Young Hae Choi, Leo FM Marcelis, Julian C Verdonk

## Abstract

Vertical farming (VF) has unparalleled capacity to highly customize plant growth environments. In VF, red and blue LED lights are predominantly used as the two main wavelengths for photosynthesis. For many plants, red light increases biomass, and blue light can increase nutritional content. Because red light is more cost-and energy-efficient to produce than blue light, refined growth recipes are imperative to mutualistically improve efficiency with crop yield and quality. This study’s aim was to balance lighting energy-use with growth and nutritional quality by using “dynamic lighting” recipes to reduce durations of high-intensity blue light. Lettuce (*Lactuca sativa* L.) was grown for 21 days at 220 μmol m^-2^ s^-1^, receiving one of five R:B ratios (R:B_100:0_, R:B_95:5_, R:B_89:11_, R:B_50:50_, and R:B_0:100_) for either the whole 18-hour photoperiod (Whole Day), the first six hours of the photoperiod (Morning), or the last six hours of the photoperiod (Evening). Morning and Evening treatments received low blue (R:B_89:11_) for the remaining 12 hours of the day. The Morning and Evening high blue treatments had greater fresh weight and leaf area than their respective Whole Day treatments, attributed to reduced instantaneous leaf photosynthesis under high blue. High blue reduced photosynthesis during only the six hours of Morning and Evening treatments, compared to the full impact of static high blue for 18-hour Whole Day treatments. Intriguingly, with only six hours of R:B_0:100_, Morning and Evening treatments had the same high anthocyanin content as lettuce grown for 18 hours under R:B_0:100_. Therefore, daily blue light fraction can be reduced by using dynamic treatments to more efficiently promote growth and nutritional quality.

## 1 Introduction

Light recipes designed to optimize plant growth are crucial for plant production in controlled environment agriculture (CEA) systems. When cultivating crops in CEA, many abiotic factors that affect plant growth can be controlled and fine-tuned, including light (both quantity/intensity and quality/spectra), nutrients, humidity, and temperature (Neo et al., 2022). One form of CEA is rising in popularity for sustainable plant production, vertical farming (VF), primarily characterized by vertically stacking layers of crop growth to reduce land usage. Plants in VF can thus be reproducibly grown more optimally with conditions designed to promote leaf growth, fruit production, or nutritional compound production, simultaneously negating effects of external factors such as herbivory, global location, and seasonal time (SharathKumar et al., 2020; van Delden et al., 2021).

Enticing as it is, VF production comes with considerable costs. The current costs required to create and run a vertical farm are high, with a considerable portion of VF expenses due to energy costs from lighting production (Banerjee and Adenaeuer, 2014; Butturini and Marcelis, 2020). Light-emitting diodes (LEDs) are often used in CEA, due to their ability to produce customizable spectra, and their increased energy efficiency and decreased heat production compared to outdated lighting methods. In VF, LEDs are commonly used to grow leafy greens with short cultivation cycles, such as lettuce, as they are more cost efficient to produce than when LEDs are used for grains (Pattison et al., 2018). In production, the most commonly used wavelengths are those most used by plant photosynthetic systems: blue light (B; 400-500 nm), and red light (R; 600-700 nm). Plants respond to these light spectra through a suite of photoreceptors, including phytochromes (Shinomura et al., 1996; Smith and Whitelam, 1997), phototropins (Christie, 2007; Inoue et al., 2008), and cryptochromes (Pedmale et al., 2016; Yu et al., 2010). Phytochromes primarily respond to red and far-red light (FR; 700-800 nm), phototropins respond to B, and cryptochromes respond mainly to B light, but also to UV-A (Assmann et al., 1985; Sharrock, 2008; Yu et al., 2010; Zeiger, 1984; Zhang et al., 2019). Through these receptors, plants respond drastically to light changes, and LED usage in agriculture and horticulture has quickly become implemented to exploit these responses in order to control growth conditions that benefit production (Pattison et al., 2018). In fact, plants grown under combined R and B LEDs have been shown to produce more plant pigments (Van Brenk et al., 2024) and increase crop yield at half of the incident energy when compared to broad-spectrum fluorescent lights (Cammarisano et al., 2021). The introduction of programmable LED modules further cements their position as ideal light sources for the highly customizable environments of VF.

Although plant responses to light can be species-or cultivar-dependent (Liu & van Iersel, 2022), plants have some common physiological responses to R and B. For example, high proportions of R are often used for cultivation as R promotes leaf expansion, is highly photosynthetically efficient, and has greater efficacy to produce the same number of photons per unit of electrical energy compared to B light (Kusuma et al., 2022, 2020). Importantly, under monochromatic R, plants exhibit a “red light syndrome”, with negatively impacted photosynthesis, stomatal function, and poor morphological characteristics such as elongated petioles and low leaf mass area (Hogewoning et al., 2010; Trouwborst et al., 2016). On the other side of the spectrum, B helps to induce photosynthesis through inducing stomatal opening (Barillot et al., 2021; Zeiger, 1984). Although B is a valuable energy source for photosynthesis, it is also a high energy source, with higher frequency than R (Consentino et al., 2015). This can stress the plant and cause the production of reactive oxygen species (ROS; Consentino et al., 2015; El-Esawi et al., 2017) and decrease photosynthesis (Liu and van Iersel, 2021). Plants may respond to B light by promoting the production of protective pigments, some that are associated with increased nutritional value and improved shelf life, such as carotenoids, flavonoids and anthocyanins (Liu et al., 2022; Min et al., 2021; Pennisi et al., 2019; Zhang et al., 2013). Carotenoids, flavonoids, and anthocyanins have high antioxidant capacity and can prevent damage from free radicals by scavenging ROS (Falcone Ferreyra et al., 2012; Khoo et al., 2017, 2011; Panche et al., 2016; Rabino and Mancinelli, 1986). Furthermore, pigments like anthocyanins can act as a “sunscreen” in cell vacuoles to filter light, diverting negative repercussions on photosynthetic machinery (Gould, 2004). Still, monochromatic B during cultivation can stunt plant growth, possibly due to hampered photosynthesis or reduced leaf expansion (Liu and van Iersel, 2021).

Conversely, in some cases, monochromatic B can cause plant elongation (Kong and Zheng, 2023). These negative or inconsistent effects of using solely R or B are alleviated by using a high R light background with low B fraction (often <15%), which restores normal function and morphology (Miao et al., 2019). R and B LEDs are therefore often used in combination, as both wavelengths are required to avoid detrimental effects of monochromatic light on plant growth (Hoenecke et al., 1992; Hogewoning et al., 2010; Trouwborst et al., 2016); VF growers using static R:B LED lighting often cultivate crops with a combination of high R (70-95%) and low B (5-30%; (Kusuma et al., 2020).

Interestingly, although programmable, customizable, LEDs have risen in popularity and accessibility, their ability to modulate their output is rarely harnessed in production and research. Often, producers select a static lighting recipe—usually with high R and low B—that meets production demands and is kept constant throughout cultivation (Lin et al., 2013; Pennisi et al., 2019; Zhang et al., 2019). This approach is inefficient as it does not consider the untapped potential from creating dynamic growth recipes to take advantage of the benefits from individual wavelengths (Kaiser et al., 2024). As stated, high proportions of R are used for more efficient energy use to increase plant growth, but high B can improve nutritional value, promoting the production of plant pigments including anthocyanins, chlorophyll, and carotenoids (Chen et al., 2021; Min et al., 2021; Van Brenk et al., 2024), which are altogether important for the grower and the target market. To simultaneously produce high R light (high R:B ratio) and high B light (low R:B) is impossible, but exposure to both environments is made possible during one photoperiod if these ratios are used dynamically at different periods of the day.

In fact, the relatively unexplored field of using a dynamic application of certain spectra at different times of the photoperiod may even be more advantageous for plant production (Kaiser et al., 2024), as a changing environment is one that plants are accustomed to (and even thrive in), having evolved by growing and responding to daily changing environmental conditions (Martínez-Vilalta et al., 2016; Ruberti et al., 2012). By applying specific light treatments such as using R:B ratios designed to drive desired plant responses at certain periods of the photoperiod, plant growth and nutrition may be increased, along with a reduction in electricity demands (Kaiser et al., 2024).

It was hypothesized that static application of low R:B (high B fraction) would increase pigments and stunt growth, as reported in Van Brenk et al. (2024). This stunted growth could be alleviated when limiting low R:B to a shorter duration of exposure, by using dynamic treatments with both periods of high R:B and periods of low R:B ratios in the photoperiod. The high R:B would save energy and benefit photosynthesis, whereas low R:B would contribute to improving quality through pigment production. The objective of this study was to identify how different exposures of R:B (high B, ≥50%, and low B, ≤11%) can be reduced in duration using dynamic spectral light treatments during the photoperiod, to improve or maintain morphological, pigment, and photosynthetic characteristics of lettuce (*Lactuca sativa*). Lettuce was grown under five different R:B light ratios at different periods of the day (morning, evening, or for the whole day), to also explore possible diel effects.

## 2 Materials and methods

### 2.1 Sowing and germination

A green-leafed multileaf lettuce, *L. sativa* cv. ‘Greenflash’, and a red-leafed multileaf lettuce, *L. sativa* cv. ‘Redflash’ (Nunhems BV, Nunhem, The Netherlands), were used for all experiments. Lettuce seeds were sown in individual rockwool plugs (Grodan, Roermond, The Netherlands), covered with a thin layer of vermiculite, and placed in a tray with tap water to imbibe the seeds and plugs. To maintain humidity, trays were covered with transparent lids. Seeds were kept in darkness for stratification (4 °C, 72 h), then transferred to a climate room to germinate. Seeds were germinated for seven days until the seedling stage, under an 18 h photoperiod at R:B_89:11_ (130 μmol m^-2^ s^-1^ PAR) and a dark period of 6 h. The B peak was ∼450 nm and the R peak was ∼655 nm, produced by GreenPower Dynamic LED modules (GPL PM 168 DRBWFR L120 G3.0 C4 N4; Signify, Eindhoven, The Netherlands). F-clean ETFE film (AGC Chemicals Europe Commercial Centre, Amsterdam, The Netherlands) was suspended beneath the LEDs for improved light distribution. The temperature was maintained at 22/21 °C day/night, the relative humidity was 65%, and CO_2_ was ambient. On the seventh day, seedlings with two fully unfurled cotyledons were selected for uniform size and quality (i.e. limited hypocotyl elongation, undamaged cotyledons). That day, ten days after sowing, seedlings in plugs were transplanted into Grodan Delta rockwool blocks (7.5 cm x 7.5 cm x 6.5 cm) previously soaked in nutrient solution [as used in Van Brenk et al. (2024)]: EC = 2.3 dS m^-1^; pH = 6-6.5; containing 12.92 mM NO_3_^−^, 8.82 mM K^+^, 4.22 mM Ca^2+^, 1.53 mM Cl^−^, 1.53 mM SO ^2−^, 1.53 mM H_2_PO_4_^−^, 1.15 mM Mg^2+^, 0.38 mM NH ^+^, 0.38 mM SiO_3_^2−^, 0.12 mM HCO_3_^−^, 38.33 μM B, 30.67 μM Fe_3_^+^, 3.83 μM Mn_2_^+^, 3.83 μM Zn^2+^, 0.77 μM Cu^2+^, and 0.38 μM Mo). After transplanting, plants were irrigated twice daily during the photoperiod, using an ebb-and-flow system that was replenished with freshly made nutrient solution approximately every four days. The transplanted seedlings in rockwool blocks were moved to growth compartments for the individual light treatments, these compartments were separated with plastic to minimize intercompartmental light contamination. Redflash and Greenflash were grown in alternating rows in each compartment.

### 2.2 Light treatments

There were 15 different light treatments, made up of two factors, red:blue ratio (R:B*_X_*) and treatment period (Morning, Whole Day, or Evening). For the first factor, the five R:B*_X_* ratios were: R:B_100:0_, R:B_95:5_, R:B_89:11_, R:B_50:50_, and R:B_0:100_ (220 μmol m^-2^ s^-1^ PAR, 18/6 h day/night). Light was provided by the GreenPower Dynamic LED modules used for germination (GPL PM 168 DRBWFR L120 G3.0 C4 N4; Signify, Eindhoven, The Netherlands). Phytochrome photostationary state (PSS) was calculated for each R:B*_X_* according to (Sager et al., 1988). For the second factor, plants received a specific R:B*_X_* spectra during one of three periods: “Morning”, “Whole day”, or “Evening” (**Figure 1**). Plants received one of the five R:B*_X_* treatments during the first six hours of the photoperiod (Morning), the final six hours of the photoperiod (Evening), or the entire 18-hour photoperiod (Whole Day). For the remaining 12 hours of Morning and Evening treatments, plants received a control ratio of R:B_89:11_. Plants were harvested at the babyleaf stage, 21 days after transplant (DAT).

**Figure 1.**
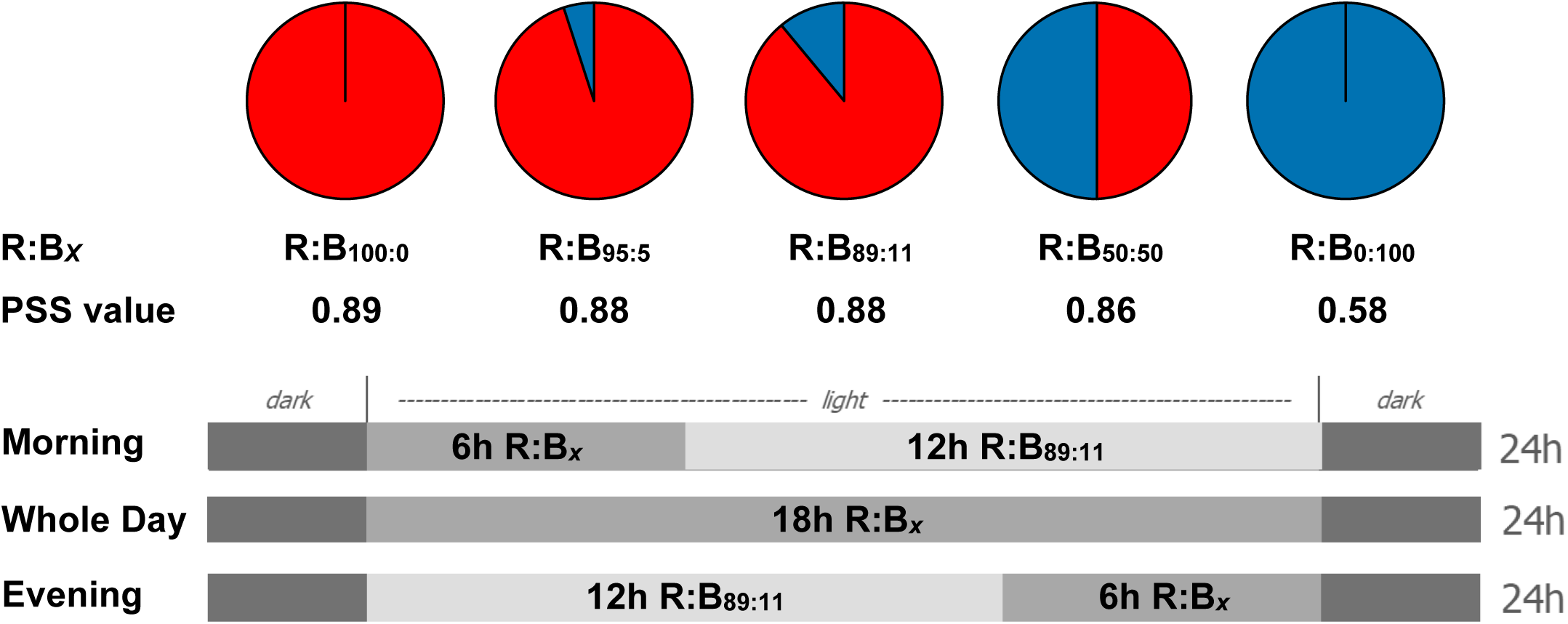
Treatment ratios and treatment periods for dynamic daily spectra Top: The specific treatment ratios of red:blue light (R:B*_X_*) and their respective phytochrome photostationary state (PSS) values. PSS was calculated for each R:B*_X_* treatment at 220 μmol m^-2^ s^-1^ PAR, according to (Sager et al., 1988). R:B*_X_* treatments only contained different fractions of blue (400- 500 nm) and red light (600-700 nm). Neither green-yellow light (500-600 nm) nor far-red light (700- 800 nm) were in the spectra. **Bottom:** The different treatment periods used in this study. The above R:B*_X_* light treatments were applied for the first six hours of the photoperiod (Morning), the Whole Day (all 18 hours), or during the last six hours of the photoperiod (Evening). When not receiving the specific R:B*_X_* treatment, Morning and Evening received R:B_89:11_.

### 2.3 Photosynthesis measurements

Photosynthesis measurements were performed during the two days preceding harvests (19 DAT). Measurements of net CO_2_ assimilation rate (A; μmol m^-2^ s^-1^) and stomatal conductance (gsw; mol m^-2^ s^-1^) were taken with a portable photosynthesis machine (Li-Cor LI-6800, Li-Cor Biosciences, Lincoln, Nebraska, USA), using a leaf chamber fluorometer with a measurement area of 2 cm^2^. The machine’s programmed conditions replicated the plant growth conditions of the climate room and were kept constant (220 µmol m^-2^ s^-1^ PPFD, 450 ppm CO_2_, 22 °C, 300 µmol s^-1^ flow rate, 10 000 RPM fan speed, and 65% relative humidity). Plants were measured twice in a day, once during the first six hours of the day (AM) and once during the last six hours of the day (PM). The measurement spectra mimicked treatment spectra during the time of measurement, with similar peak wavelengths between the measurement and growth light conditions. For example, a Morning R:B_50:50_ plant was measured under R:B_50:50_ during the AM measurement and at R:B_89:11_ during the PM measurement. Either the first or second true leaves were measured, with priority for the second leaf, but reliant on minimum leaf size limitations of the measurement chamber. Leaves fully covered the leaf chamber gasket. After allowing plants to stabilize for at least 5 minutes, the machine measured A and gsw for 4 seconds, followed by a 10-second delay, then measured again for 4 seconds; then, these two 4-second measurements were averaged for a singular value for either A or gsw.

### 2.4 Stomatal imprinting

Stomatal imprints were performed during the two days preceding harvests (19 DAT). Silicone impressions were used to create a negative imprint of leaf stomata from the first or second fully developed leaves, as in Geisler et al. (2000) and Zhang et al. (2022). From two plants per treatment, three imprints on both the adaxial and abaxial sides of leaves were collected using dental silicone (Zhermack elite HD+ light body a-silicone, Zhermack SpA, Rovigo, Italy), which was removed when set. These silicone imprints were covered with clear acrylic nail polish; after the polish dried, it was removed from the chip and placed on a microscope slide, as a positive imprint of stomata. The clear acrylic prints were then magnified 80x using an optical microscope (Leitz Aristoplan 020-503.030, Ernst Leitz GmbH, Rijswijk, Netherlands) and visualized digitally with a camera (Axiocam 305 colour, Carl Zeiss Microscopy GmbH, München, Germany). Each leaf side for each treatment was photographed in four different areas of the imprint. Images were collected via Zen 3.3 software (Blue edition version 3.3.89, Carl Zeiss Microscopy GmbH, München, Germany). The resulting images were analyzed by manually counting stomata using ImageJ (National Institutes of Health, Bethesda, Maryland, USA). Stomatal density was measured by dividing the number of stomata counted in the total photographed area of known size (1080 μm x 900.72 μm), converting to stomata mm^-2^.

### 2.5 Morphological measurements

Photographs were taken of four representative plants, per treatment, at each harvest using a digital camera (Nikon D7200, Nikon, Tokyo, Japan) and tripod. Photographs were taken from above and from the side of the lettuce. For each harvest, all overhead photos were taken with the same manual shutter speed, ISO, aperture, and zoom, with the same lens, performed in the same windowless room, under the same light conditions. Photographs from the side were also performed with the exact same settings as other side-view photos. The only difference in camera settings was the ISO when photographing Redflash or Greenflash, as the lighter colouration of Greenflash reflected more light, compensated by reducing the ISO.

Immediately after being photographed, 16 whole lettuce plants per treatment, per cultivar, were harvested by cutting above the root below the cotyledons. Each plant was measured for fresh weight with an analytical balance (DK-6200-C-M, NL-220-C-M, AllScales Europe, Veen, The Netherlands), leaf count (number of leaves taller than 1 cm, excluding cotyledons), and leaf area (Li-Cor LI-3100C, Li-Cor Biosciences, Lincoln, Nebraska, USA). Following harvest, 12 of the 16 plants were dried in a forced-air oven (Elbanton Special Products by Hettich Benelux, Geldermalsen, The Netherlands) for three days (70 °C for 24 h, then 100 °C for 48 h) and weighed for dry weight. Specific leaf area (SLA) of each plant was calculated using dry weight and leaf area data. The four remaining plants per treatment were flash-frozen in liquid nitrogen and stored at-80 °C, to be used for pigment and metabolite analysis. These procedures were repeated for all four replicate experiments (*n* = 4).

### 2.6 Pigment and metabolite analysis

The four lettuce plants per treatment that were frozen and stored at-80 °C were freeze-dried for 72 h in a freeze-dryer (Edwards Modulyo II, Edwards High Vacuum Int., Sussex, United Kingdom). The freeze-dried samples of whole plants were individually ground to a powder using a ball mill (MM 400, Retsch, Dale i Sunnfjord, Norway), then equal weights of each of the four plants were combined as a pooled sample; this pooling was performed for each replicate experiment, per treatment ratio, per treatment period, per cultivar.

Chlorophyll a, chlorophyll b, carotenoid, and total flavonoid concentrations were determined using ethanolic extraction of ∼5 mg of the above-described ground and pooled freeze-dried tissue, according to relevant sections of the rainbow protocol (López-Hidalgo et al., 2021), with some modifications for diluting the final measured photosynthetic pigment sample 1:4. The exact weight of each sample was determined using a high accuracy balance (AT21, Mettler-Toledo, Columbus, Ohio, USA). The absorbance of the final ethanolic extraction in the wells of a 96-well microplate (Cellstar® sterile F-bottom, Greiner Bio-One B.V., Alphen aan den Rijn, The Netherlands) was quantified utilizing a SpectraMax iD3 microplate reader (Molecular Devices; San Jose, CA, United States). Total flavonoids were measured at 415 nm and expressed as mg of quercetin equivalents per gram dry weight, compared to a quercetin standard curve (from 0 to 1 mg/mL). Chlorophyll a, chlorophyll b, and carotenoids were determined by measuring the absorbance at 664, 649, and 470 nm, then calculated using the following equations from Lichtenthaler and Wellburn (1983):

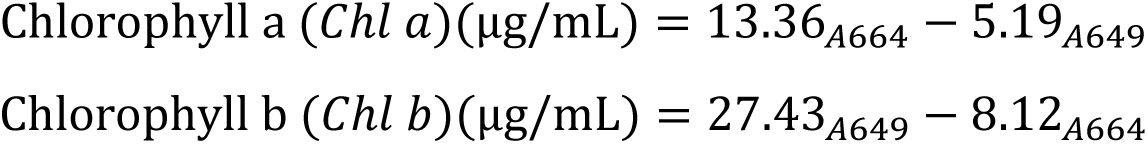

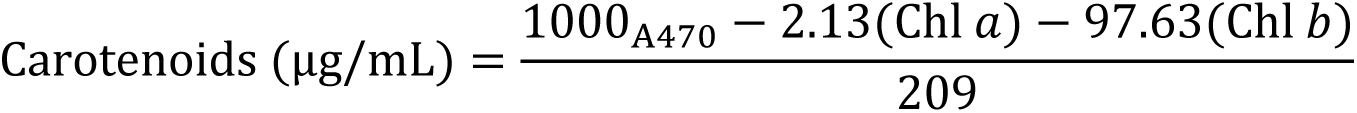

Relative anthocyanin concentration was determined based on Neff & Chory (1998) with some modifications, using 5 mg of pooled freeze-dried tissue. Briefly, samples were mixed with 300 μL methanol and 1% hydrochloric acid (HCl), extracted overnight in the dark (4 °C). The next day, 200 μL ddH_2_O was added, followed by 500 μL chloroform. After briefly vortexing the tubes three times, samples were spun in a centrifuge (15000 x g, 3 min, 22 °C). A dilution step was added, using 100 μL of the aqueous solution, combined with 900 μL methanol with 1% HCl. Then, 150 μL of the diluted solution was pipetted into three different wells of a microplate. To determine anthocyanin concentration, the absorbances at 530 and 657 nm were measured for each well of a microplate and quantified utilizing a SpectraMax iD3 microplate reader, compared to a blank containing methanol and 1% HCl. Total relative anthocyanin concentration was calculated as relative units of (A_530_-A_657_) per gram dry weight.

### 2.7 ^1^H NMR analysis

30 mg of ground and freeze-dried lettuce sample material was placed into 2 mL microtubes, performed using two plants per treatment, per replicate experiment. Each tube was mixed with 1 mL of CD_3_OD (deuterated methanol), CD_3_OD containing 0.418 mM HMDSO (hexamethyldisiloxane), or a CD_3_OD- KH_2_PO_4_ buffer (pH 6.0, with 0.58 mM TMSP-d_4_ in a 1:1, v/v ratio). The tubes were vortexed (1 min, 22 °C) to ensure proper mixing. To further enhance extraction, the microtubes were subjected to ultrasonication (20 min, 22 °C). Following this, the samples were centrifuged (12000 x g, 10 min, 22 °C) to obtain a clear supernatant, from which 300 μL was transferred into a 3 mm NMR tube.

NMR measurements were conducted using a Bruker Avance-III 600 MHz standard bore liquid-state NMR spectrometer (Bruker, Billerica, Massachusetts, USA) operating at a magnetic field strength of 14.1 Tesla, with the ^1^H nucleus resonating at 600.13 MHz. The spectrometer was equipped with a TCI cryoprobe optimized for H&F/C/N-D detection and featuring a Z-gradient. Experiments utilized 3 mm NMR tubes sourced from Cortecnet (Les Ulis, France). The temperature was maintained at a constant 298 K throughout the measurements, and CH_3_OH-d_4_ was employed as an internal lock. For each proton experiment, a 30-degree pulse of 2.64 milliseconds duration was applied at a power level of 5.5 W, resulting in a free induction decay (FID) resolution of 0.36 Hz. A total of 64 scans were performed with a relaxation delay of 1.5 seconds and an acquisition time of 2.7 seconds, leading to a total experiment duration of approximately 5 minutes. The suppression of the water signal was achieved using a pre-saturation method with low-power selective irradiation at 0.3 Hz targeting H_2_O at 4.87 ppm. The collected time-domain data was converted to the frequency domain through Fourier transformation, utilizing an exponential window function with a line broadening parameter of 0.3 Hz for smoothing. The resulting spectra were manually phased, baseline corrected and calibrated to reference standards: TMSP-d4 at 0.0 ppm or HMDSO at 0.06 ppm.

### 2.8 Multivariate and statistical analysis of ^1^H NMR data

SIMCA-P software (version 18.0.1, Sartorius, Amersfoort, The Netherlands) was utilized to perform multivariate data analysis based on matrices derived from ^1^H NMR data. The bucketed dataset was analysed by principal component analysis (PCA) and orthogonal projections to latent structures discriminant analysis (OPLS-DA) using PCs reduced by PCA. In addition, to assess the clustering of growth conditions on sample metabolic variation, a soft independent model of class analogy (SIMCA) analysis was performed using lettuce samples as PCA classes separately in each blue light condition.

### 2.9 Statistical design and analysis

Four replicate experiments were conducted, representing four blocks (*n* = 4; 16 plants per treatment), with each combined treatment period and R:B*_X_*. For each experiment, treatments were randomly assigned to different growth compartments. Analysis for significance was completed using two-way analysis of variance (ANOVA) in blocks, with both cultivars analyzed separately. For morphology (using 16 plants per treatment) and metabolites (from pooled tissue of four randomly selected plants per treatment), the factors for the two-way ANOVA were the blue light % in a R:B background and the treatment period. For this, *P*_Blue_ was the probability of an effect due to R:B*_X_* blue content; *P*_Period_ was the probability of an effect due to the treatment period of Morning, Whole Day, or Evening; and *P*_int_ was the probability of an interactive effect between R:B*_X_* blue content and treatment period (all using *a* = 0.05). For photosynthesis measurements (assimilation and stomatal conductance; from three randomly selected plants per treatment) the factors were the blue light % in a R:B background and the time of measurement. These photosynthesis data were analyzed using a split-plot design with the whole plots being the treatment period (Morning, Whole Day, Evening) and the sub-plots being the time of measurement (AM or PM measurement time). For these, *P*_Blue_ was the probability of an effect due to R:B*_X_* blue content; *P*_AM/PM_ was the probability of an effect due to the time of measurement; and *P*_Int_ was the probability of an interactive effect between R:B*_X_* blue content and measurement time (all using *a* = 0.05). Finally, stomatal density was determined using two randomly selected plants for two replicate experiments (*n* = 2), the factors for the two-way ANOVA were blue light % in a R:B background and the treatment period. Following each ANOVA test (*p* < 0.05), multiple comparison analysis was performed with Fisher’s unprotected least significant difference (LSD) tests (*p* < 0.05). When creating figures considering total daily B fraction, a trendline for all treatment periods was created if 95% of variance could be accounted for with a polynomial trendline. Otherwise, individual trendlines were produced for each treatment period, using an ANOVA test to check for a linear or quadratic effect of B fraction (*a* = 0.05). Due to facility accessibility limitations, the Morning treatment periods were not performed at the same time as the Whole Day and Evening treatment periods. To address this discrepancy properly for statistic analysis, the control treatment of the Morning R:B_89:11_ was used to scale to the Whole Day and evening R:B_89:11_ data, per replication block. This scaling factor was applied to each treatment within a block, resulting in the scaling of all Morning treatment data to those of the Whole Day and Evening R:B_89:11_ treatment data.

## 3 Results

### 3.1 Plant growth reduction by B light was improved by using diurnal treatments

Overall, for both cultivars, fresh weight decreased with increased B for Whole day, Morning and Evening treatments (**Figure 2A; Figure 2B**). The effects were stronger for Whole Day treatments than both Morning and Evening treatments, which were overall the same for each individual R:B*_X_* ratio. However, when plotted versus the daily B fraction (averaged over the entire photoperiod), plant growth corresponded to the daily B fraction, independent of whether B was given in the Morning, Whole Day, or Evening—except for the Morning and Evening R:B_0:100_ treatments in Greenflash (**Figure 2C; Figure 2D**). For Greenflash, this short period (6h) of monochromatic B resulted in more growth than when the R:B treatments were equally spread over the Whole Day. These described patterns of fresh weight were the same as those of dry weight and leaf area, for each treatment and for both cultivars (**Supplemental Figures 1 and 2**). Contributing to these data, the leaf number for Greenflash and Redflash both decreased with increased B fraction, but this was again less impacted in the six-hour Morning and Evening treatments compared to Whole Day treatments (Supplemental Table 1).

**Figure 2.**
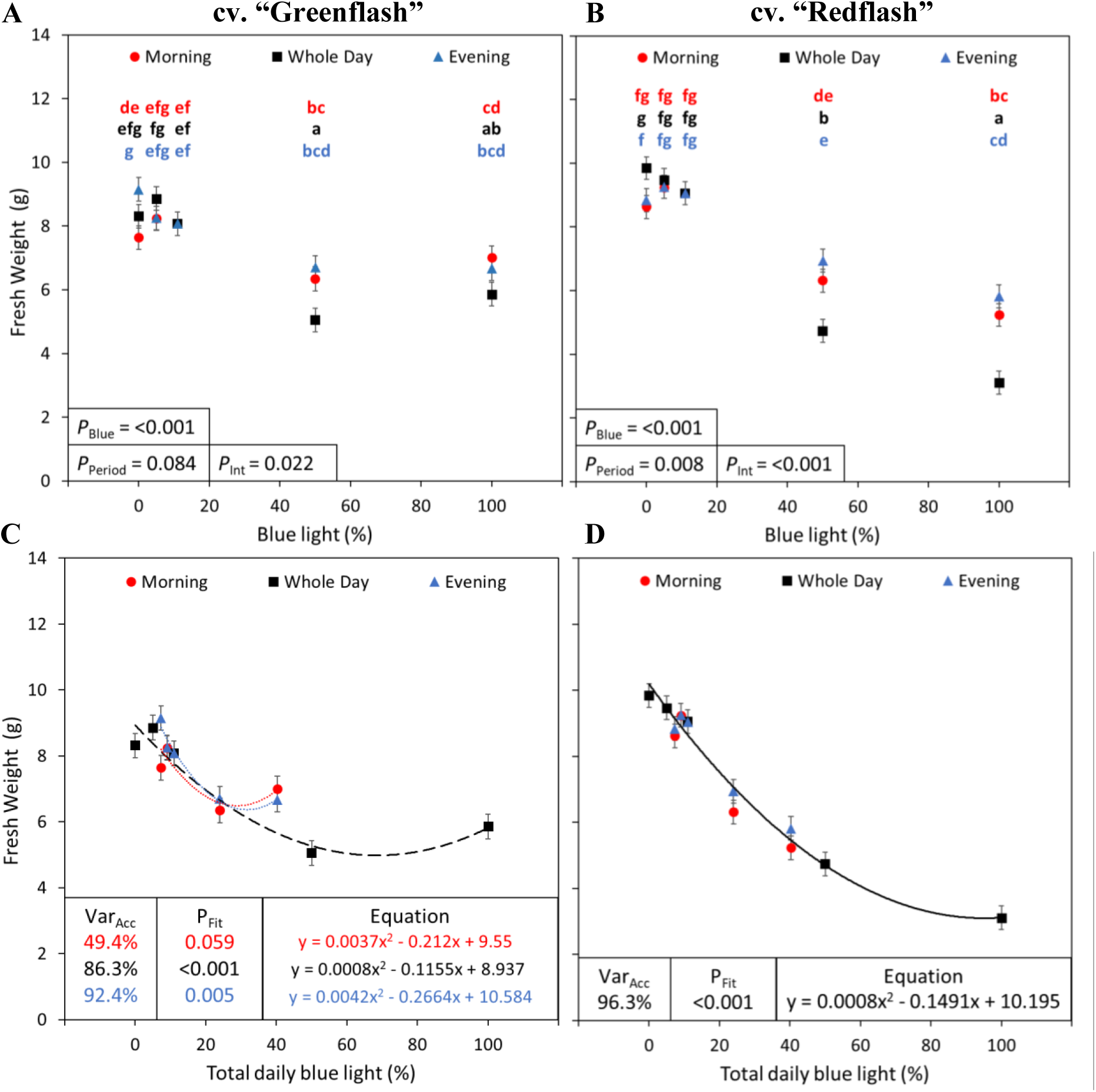
Fresh weight of two lettuce cultivars grown under five different red:blue ratios applied during three different parts of the day. Fresh weight (g) of lettuce cv. “Greenflash” (**A, C**) and cv. “Redflash” (**B, D**) grown for 21 days. Five different R:B*_X_* ratios (R:B_100:0_, R:B_95:5_, R:B_89:11_, R:B_50:50_, and R:B_0:100_) were applied either during the six Morning hours, six Evening hours, or during Whole Day for all 18 hours of the day. The fraction of B light during the treatment period (**A, B**) or total B fraction for the whole photoperiod (**C, D**) were used as X-axes. Datapoints represent means with standard error means of four growth cycles (*n* = 4), each consisting of sixteen replicate plants. For panels (**A**) and (**B**), different letters indicate significantly different values for each combination of R:B*_X_* and treatment period, according to an unprotected Fisher LSD test (*a* = 0.05). *P*_Blue_ = probability of an effect due to R:B*_X_* blue content; *P*_Period_ = probability of an effect due to the treatment period of Morning, Whole Day, or Evening; *P*_int_ = probability of an interactive effect between R:B*_X_* blue content and treatment period; Var_Acc_ = percent variance accounted for by regression line; *P*_Fit_ = probability of a linear or quadratic trend.

### 3.2 Monochromatic treatment effects on morphology were alleviated with diurnal application

The sizes of plants seen in representative photographs aligned with the plant growth metrics (**Figure 3, Supplemental Figures 3 and 4**). The most obvious phenotypic impacts occurred in plants subjected to Whole Day treatments, for both Greenflash and Redflash. Both cultivars, when grown under monochromatic R (R:B_100:0_) for the Whole Day displayed lighter-green pigmentation, elongated leaves and petioles, and increased leaf curvature compared to plants of other treatments (**Figure 3A; Figure 3B; Figure 3F; Figure 3G**). Under monochromatic B (R:B_0:100_), Greenflash appeared darker-green and had more vertical, hyponastic, leaf orientation compared to R:B_89:11_ (**Figure 3E**). Although noticeably less hyponastic than Greenflash, Redflash R:B_0:100_ plants also grew more vertically than in other treatments, their leaf colour a deep, reddish-purple hue (**Figure 3J**).

**Figure 3.**
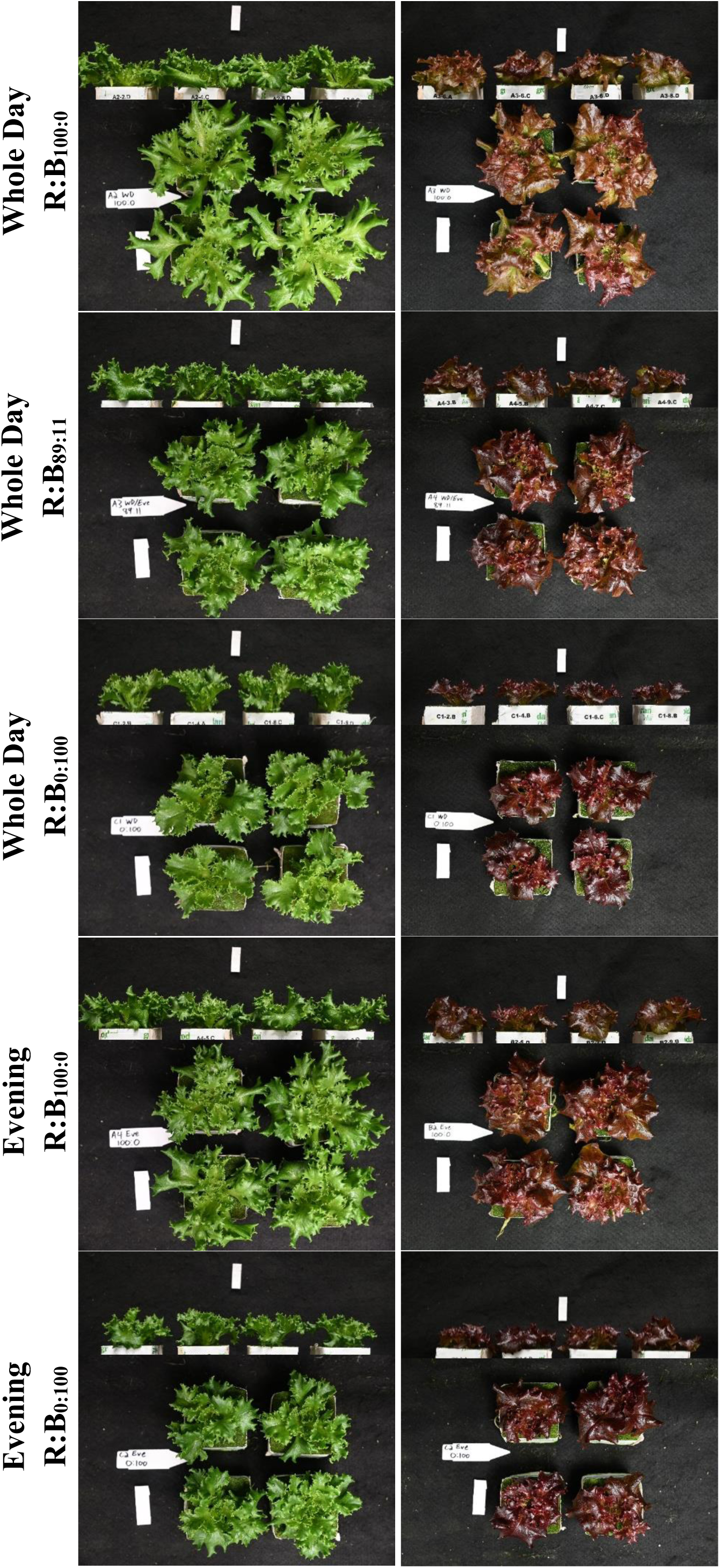
Representative photos (side and overhead views) of green lettuce cv. “Greenflash” and red lettuce cv. “Redflash” grown for twenty-one days under different R:B*_X_* treatments (**A - E**) cv. “Greenflash”, (**F - J**) cv. “Redflash”. Treatments shown here as: (**A, F**) Whole Day R:B_100:0_; (**B, G**) R:B_89:11_ as control; (**C, H**) Whole day R:B_0:100_; (**D, I**) Evening R:B_100:0_; (**E, J**) and Evening R:B_0:100_.

Notably, the Morning and Evening treatments— with their reduced treatment durations—alleviate the strong phenotypic impacts of each Whole Day treatment. This included the vertical growth and curling of leaves characteristic of plants grown under monochromatic R or B, which were not observed for Morning or Evening treatments. As expected, Morning, Whole Day, and Evening R:B_89:11_ treatments did not have noticeable phenotypic deviations, because all these plants received 18 hours of R:B_89:11_.

### 3.3 Metabolites and pigments are differentially affected by light treatments, for both cultivars

#### 3.3.1 The metabolomic profiles of spectra within treatment periods are identifiably segregated

For both cultivars, when comparing all of the Morning, Whole Day, and Evening treatments together, these three periods of R:B*_X_* application had metabolite profiles that were overall different (**Figure 4A; Figure 4E**;). These three treatment periods largely segregated into their respective groupings by their period of treatment, with some overlap (as they had one treatment in common, R:B_89:11_). When analyzing the R:B*_X_* spectra treatments within each of the three individual treatment periods, it was found that overall the low B treatments R:B_100:0_, R:B_95:5_, and R:B_89:11_ had overlapping profiles, but the high B treatments R:B_50:50_ and R:B_0:100_ had distinct separation from the low B group and from each other (**Figure 4B-D; Figure 4F-H**). This was the case for each of Morning, Whole Day, and Evening treatments, and for both cultivars; this indicates that B fraction affects the plants’ metabolic profile.

**Figure 4.**
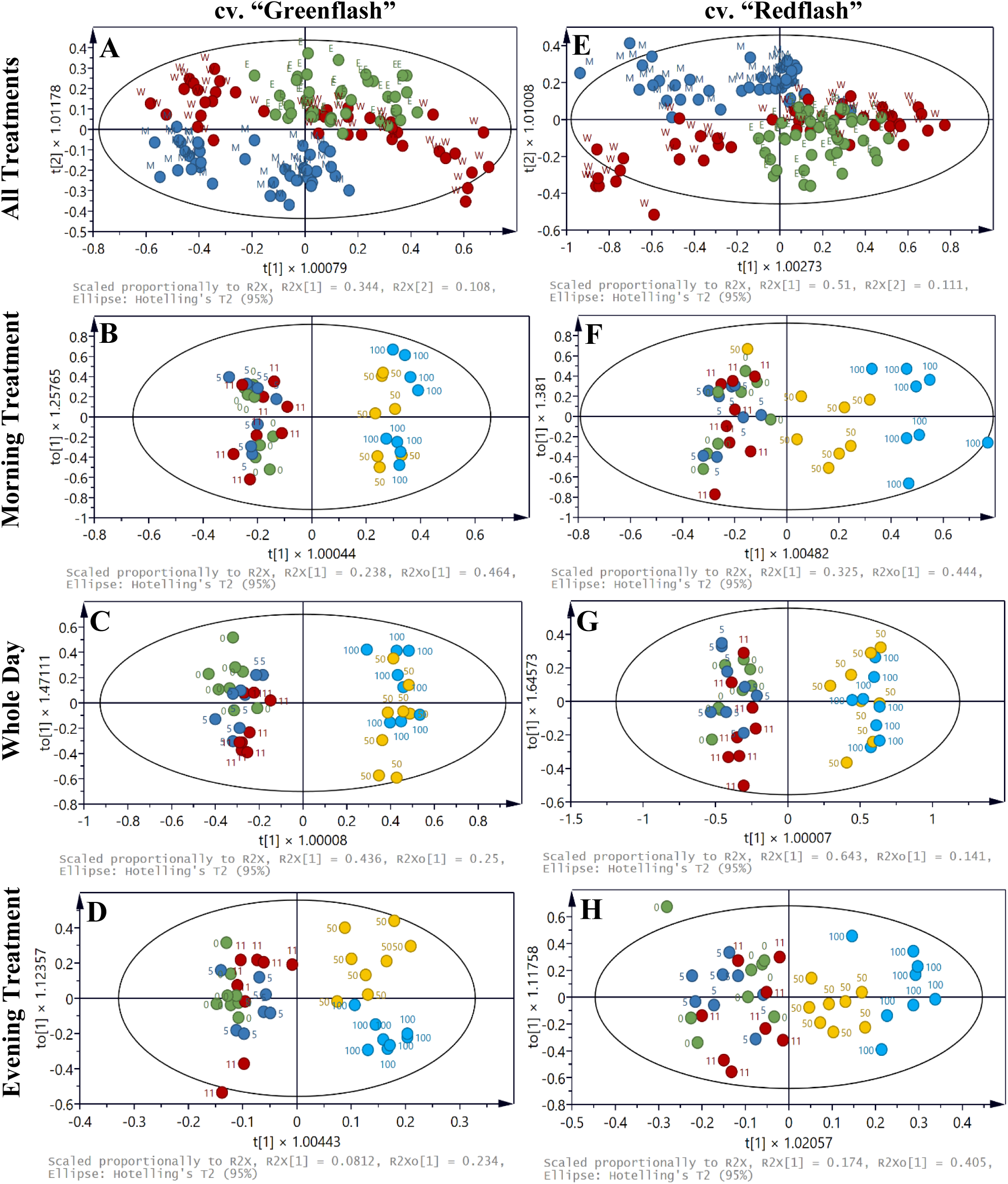
Metabolomic profiles of two lettuce cultivars grown under five different red:blue ratios applied during three different parts of the day. Orthogonal partial least squares discriminant analysis (OPLS-DA) plots of (**A - D**) green lettuce cv. “Greenflash” and (**E - H**) red lettuce cv. “Redflash” grown for 21 days. Five different R:B*_X_* ratios (R:B_100:0_, R:B_95:5_, R:B_89:11_, R:B_50:50_, and R:B_0:100_) were applied either during the six Morning hours (labelled as M in panels **A** and **E**), six Evening hours (labelled as E in panels **A** and **E**), or during Whole Day for all 18 hours of the day (labelled as W in panels **A** and **E**). For panels **B - D** and **F - H**, the R:B*_X_* ratio is labelled by its blue light fraction (0, 5, 11, 50, or 100). These plots compare the metabolomic separation between R:B*_X_* treatments for (**B, F**) Morning, (**C, G**) Whole Day, and (**D, H**) Evening treatments. Plots were generated using the relative intensities of NMR spectral bins. Multivariate data analyses identified the different classes of primary and secondary metabolites that contributed to each sample’s different metabolome profile.

#### 3.3.2 High blue light increases chlorophyll and reduces carotenoids in both cultivars

In Greenflash, the chlorophyll concentration was highest for R:B_50:50_, irrespective of the application time (**Figure 5A, Supplemental Table 1**). Six hours of Morning or Evening R:B_100:0_ treatments had higher chlorophyll than Whole Day R:B_100:0_, suggesting that having the presence of any content of blue light can improve chlorophyll levels. In Redflash, slight differences were found for high B fractions: Whole Day R:B_0:100_ had the highest chlorophyll levels (**Figure 5B, Supplemental Table 1**), which agrees with higher blue leading to higher chlorophyll. Redflash also showed a decrease in chlorophyll a/b ratio with increasing blue fraction, most obviously in the Whole Day treatments, but also seen for Morning and Evening treatments (Supplemental Table 1). For both cultivars, total carotenoids decreased for R:B_50:50_ and R:B_0:100_, most strongly for Whole Day exposure and to a lesser extent for Morning and Evening treatments (**Figure 5C; Figure 5D**). Finally, both chlorophyll and carotenoids were roughly double in Greenflash compared with Redflash (**Figure 5**).

**Figure 5.**
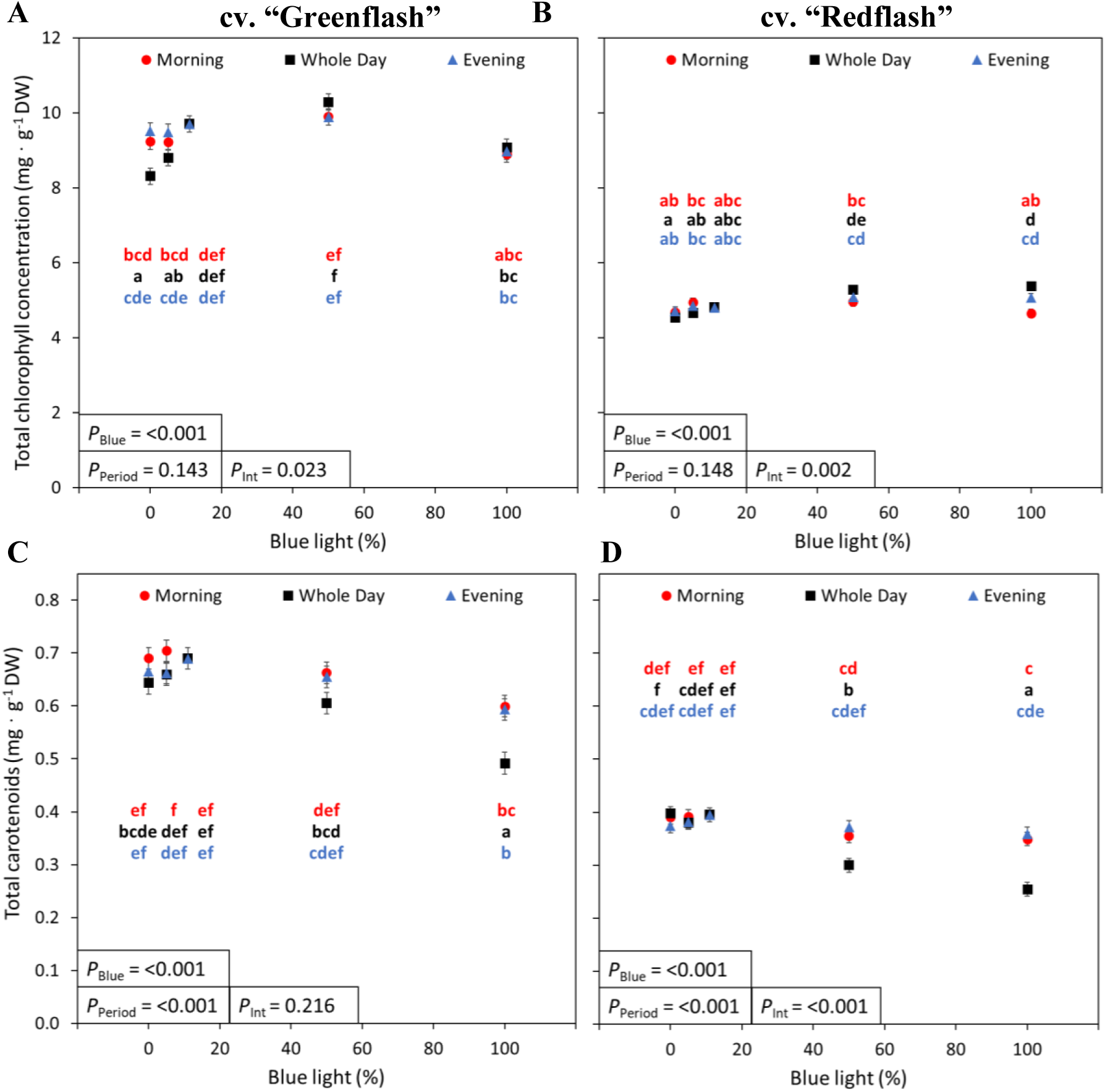
Total chlorophyll and carotenoid concentration of two lettuce cultivars grown under five different red:blue ratios applied during three different parts of the day. Total chlorophyll (**A, B**) and carotenoid (**C, D**) concentration (mg · g^-1^ DW) of lettuce cv. “Greenflash” (**A, C**) and cv. “Redflash” (**B, D**) grown for 21 days. Five different R:B*_X_* ratios (R:B_100:0_, R:B_95:5_, R:B_89:11_, R:B_50:50_, and R:B_0:100_) were applied either during the six Morning hours, six Evening hours, or during Whole Day for all 18 hours of the day. The fraction of B during the treatment period (**A, B**) or total B fraction for the whole photoperiod (**C, D**) were used as X-axes. Datapoints represent means with standard error means of four growth cycles (*n* = 4), each consisting of sixteen replicate plants. For panels (**A**) and (**B**), different letters indicate significantly different values for each combination of R:B*_X_* and treatment period, according to an unprotected Fisher LSD test (*a* = 0.05). *P*_Blue_ = probability of an effect due to R:B*_X_* blue content; *P*_Period_ = probability of an effect due to the treatment period of Morning, Whole Day, or Evening; *P*_int_ = probability of an interactive effect between R:B*_X_* blue content and treatment period; Var_Acc_ = percent variance accounted for by regression line; *P*_Fit_ = probability of a linear or quadratic trend.

#### 3.3.3 Diurnal treatments accumulate the same anthocyanin concentration as static high blue, but with less total daily blue light, similar effects with flavonoids

Anthocyanins were only found in the red cultivar, Redflash (**Figure 6, Supplemental Figure 5**). Aligning with the colouration of its leaves, the lowest anthocyanin content was from R:B_100:0_ Whole day plants. Conversely, the R:B_0:100_ plants had the deepest-coloured red leaves, corresponding with the highest anthocyanin content (**Figure 3J; Figure 6A**). Anthocyanins for Morning and Evening treatments were not significantly different for the respective R:B*_X_* treatments (i.e. Morning R:B_0:100_ = Evening R:B_0:100_), and only differed in content from Whole Day treatments in R:B_100:0_ treatments (**Figure 6A**). Interestingly, plotting the total daily B received for all treatments showed that high B in Morning and Evening treatments had higher anthocyanin accumulation than the trend of Whole Day anthocyanin content (**Figure 6B**).

**Figure 6.**
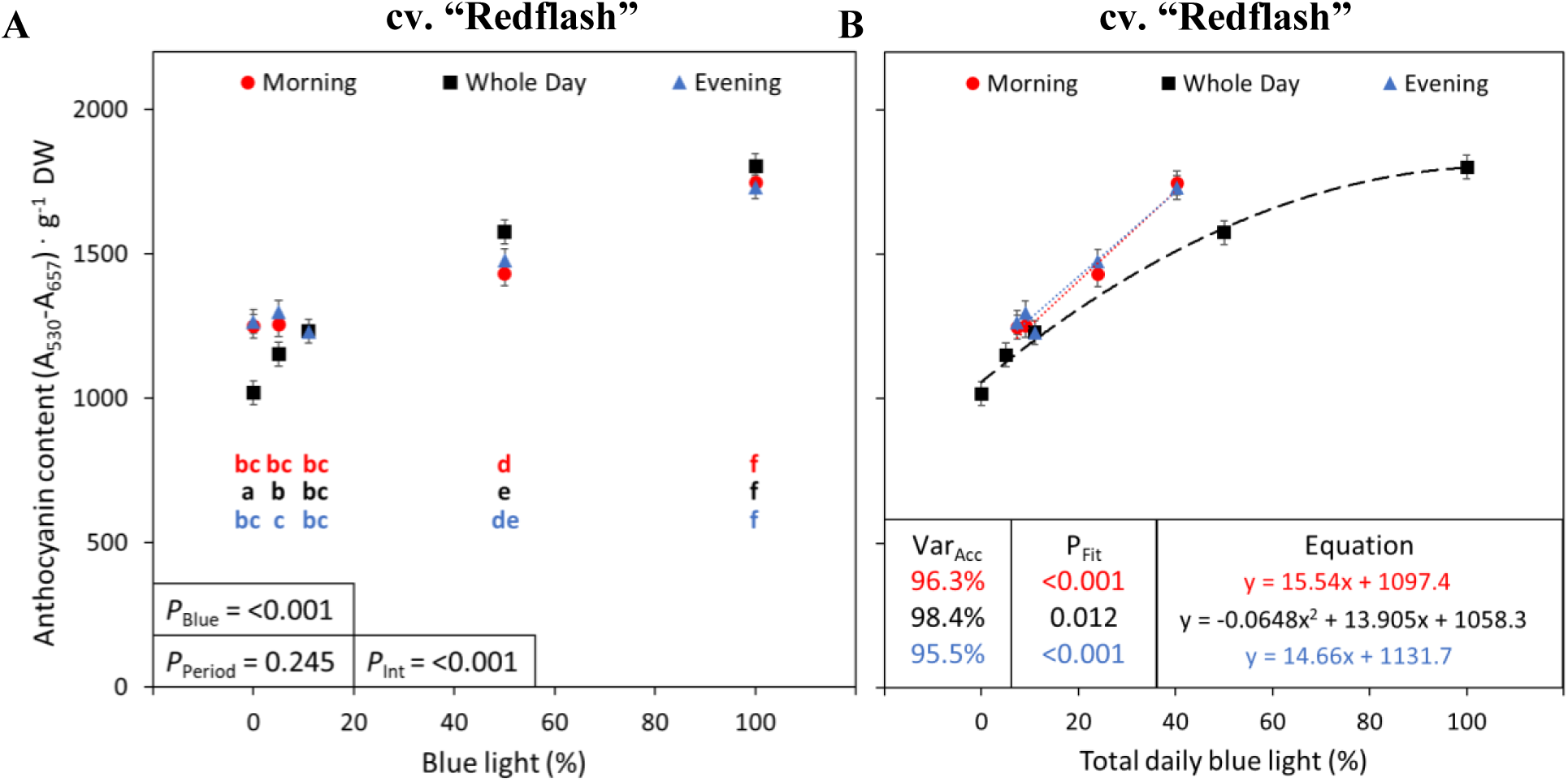
Relative anthocyanin concentration of “Redflash” lettuce grown under five different red:blue ratios applied during three different parts of the day. Relative anthocyanin concentration (A_530_-A_657_ · g^-1^ DW) of lettuce cv. “Redflash” grown for 21 days. Five different R:B*_X_* ratios (R:B_100:0_, R:B_95:5_, R:B_89:11_, R:B_50:50_, and R:B_0:100_) were applied either during the six Morning hours, six Evening hours, or during Whole Day for all 18 hours of the day. The fraction of B during the treatment period (**A, B**) or total B fraction for the whole photoperiod (**C, D**) were used as X-axes. Datapoints represent treatment means with error bars representing standard error means of four growth cycles (*n* = 4), each consisting of sixteen replicate plants. For panels (**A**) and (**B**), different letters indicate significantly different values for each combination of R:B*_X_* and treatment period, according to an unprotected Fisher LSD test (*a* = 0.05). *P*_Blue_ = probability of an effect due to R:B*_X_* blue content; *P*_Period_ = probability of an effect due to the treatment period of Morning, Whole Day, or Evening; *P*_int_ = probability of an interactive effect between R:B*_X_* blue content and treatment period; Var_Acc_ = percent variance accounted for by regression line; *P*_Fit_ = probability of a linear or quadratic trend.

The parent class of anthocyanins, the flavonoids, also increased with B fraction. For Greenflash and Redflash, the Whole day treatments had the lowest flavonoids with low B treatments and the highest flavonoids with high B treatments (**Figure 7A**; **Figure** 7**B**). Both Morning and Evening showed a less strong increase than Whole Day with the treatment B fraction (**Figure 7A**; **Figure** 7**B**).When considering total daily B fraction, for Greenflash, these aligned well with the trend for each Whole Day treatments (**Figure 7C**). Interestingly, when considering total daily B fraction, the Redflash Morning and Evening high B treatments exceeded the Whole Day trend of flavonoid content (**Figure 7D**), much like the pattern of anthocyanin content. Interestingly, the biosynthesis of anthocyanins in Redflash (but not in Greenflash) did not cause a large difference in total flavonoid content between the two cultivars, except potentially for the dynamic R:B_50:50_ and R:B_0:100_ treatments.

**Figure 7.**
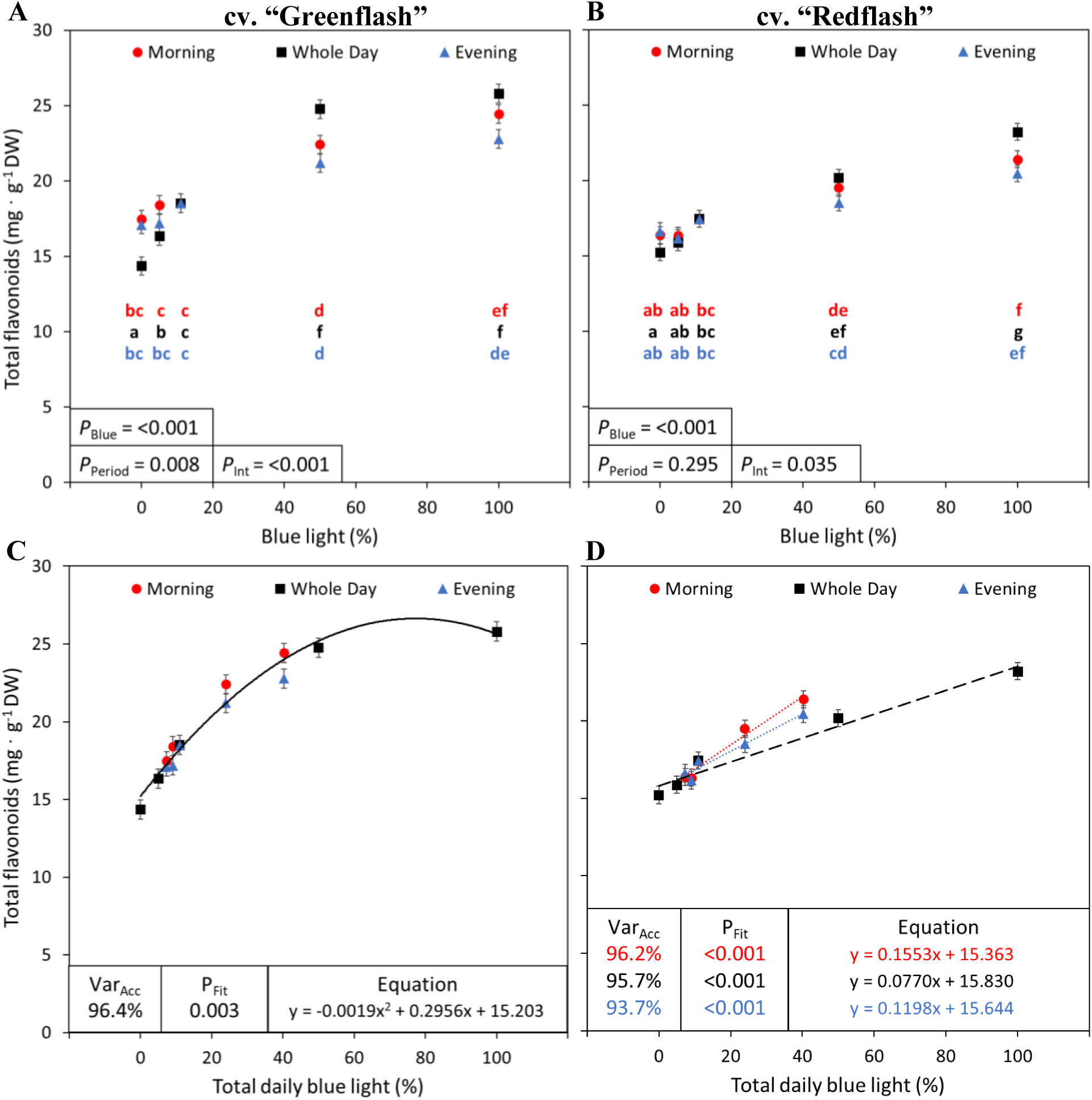
Flavonoid concentration of two lettuce cultivars grown under five different red:blue ratios applied during three different parts of the day. Flavonoid concentration of lettuce cv. “Greenflash” (**A, C**) and cv. “Redflash” (B, D) grown for 21 days. Five different R:BX ratios (R:B_100:0_, R:B_95:5_, R:B_89:11_, R:B_50:50_, and R:B_0:100_) were applied either during the six Morning hours, six Evening hours, or during Whole Day for all 18 hours of the day. The fraction of B during the treatment period (**A, B**) or total B fraction for the whole photoperiod (**C, D**) were used as X-axes. Datapoints represent treatment means with error bars representing standard error means of four growth cycles (*n* = 4), each consisting of sixteen replicate plants. For panels (**A**) and (**B**), different letters indicate significantly different values for each combination of R:BX and treatment period, according to an unprotected Fisher LSD test (*a* = 0.05). PBlue = probability of an effect due to R:B*_X_* blue content; *P*_Period_ = probability of an effect due to the treatment period of Morning, Whole Day, or Evening; *P*_int_ = probability of an interactive effect between R:B*_X_* blue content and treatment period; *Var*_Acc_ = percent variance accounted for by regression line; *P*_Fit_ = probability of a linear or quadratic trend.

### 3.4 Photosynthesis is primarily affected by the instantaneous treatment light spectra

Although Greenflash assimilation and conductance were comparatively higher than Redflash, both cultivars shared some overlapping patterns (**Figure 8** and **Figure 9**). For both cultivars, assimilation rates and stomatal conductance measured during the AM measurement time (measurements taken in the first six hours of the photoperiod) were overall higher than each respective PM measurement (measurements during the last six hours). Further, the assimilation and stomatal conductance of R:B_100:0_, R:B_95:5_, and R:B_89:11_ overall did not significantly differ from each other during AM measurements or PM measurements, respectively.

**Figure 8.**
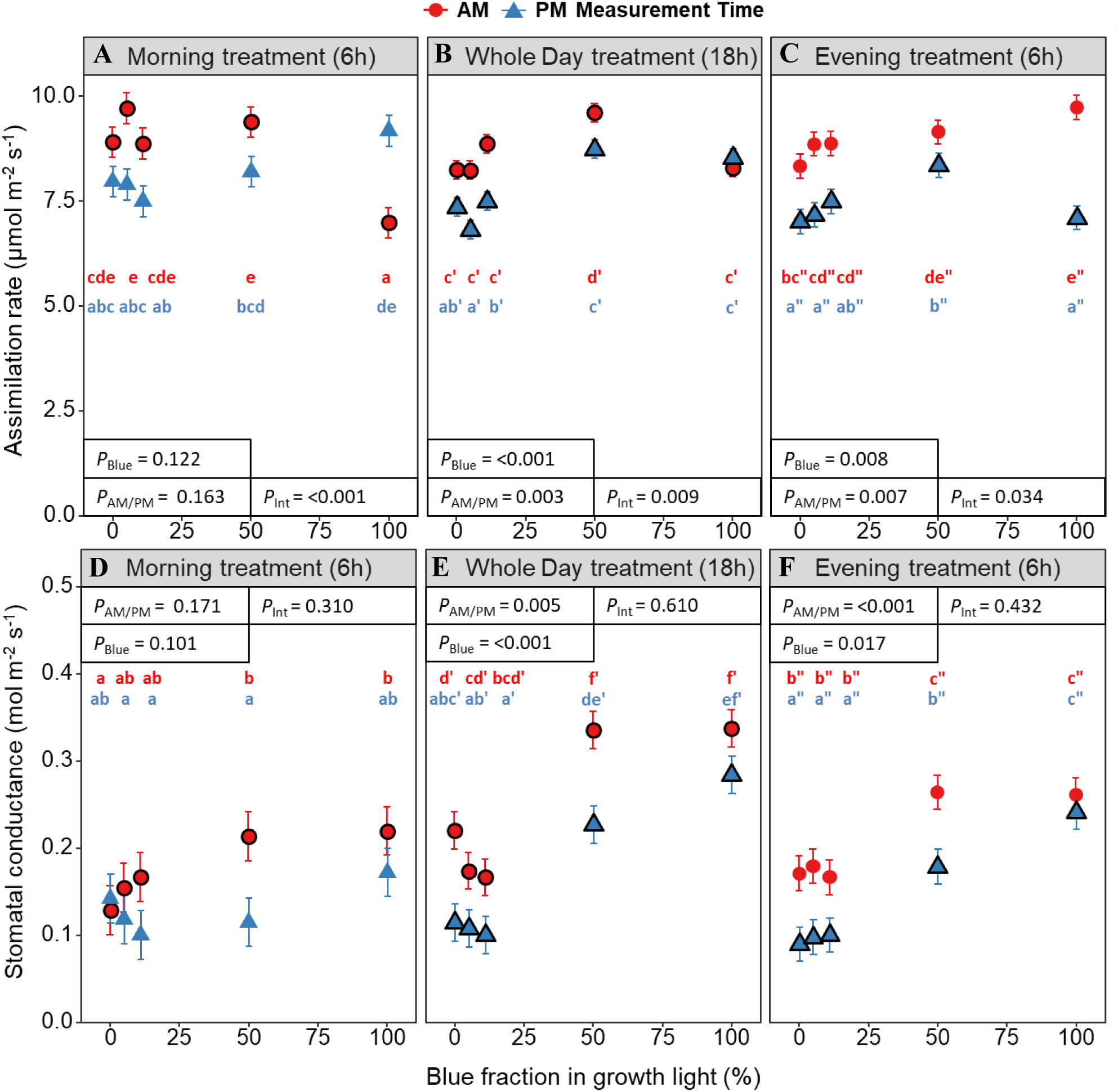
Assimilation rate and stomatal conductance of green lettuce grown under different red:blue treatments. Measurements during the Morning (AM) and Evening (PM) for assimilation rates (**A-C**; µmol CO_2_ m^−2^ s^−1^) and stomatal conductance (**D-F**; mol CO_2_ m^−2^ s^−1^) of lettuce cv. “Greenflash” grown for 21 days. Five different R:B*_X_* ratios (R:B_100:0_, R:B_95:5_, R:B_89:11_, R:B_50:50_, and R:B_0:100_) were applied either during the six Morning hours, six Evening hours, or during Whole Day for all 18 hours of the day. Shape outlines indicate plants receiving the specific R:B*_X_* treatment during measurements, otherwise plants received R:B_89:11_. Datapoints represent treatment means and error bars represent standard error means of four growth cycles (*n* = 4), each with three replicate plants. Different letters indicate significantly different values for each combination of R:B*_X_* and measurement time, according to an unprotected Fisher LSD test (*a* = 0.05). Apostrophes indicate significance per treatment, for Morning (no apostrophe), Whole Day (′), and Evening (″). *P*_Blue_ = probability of an effect due to R:B*_X_* blue content, *P*_AM/PM_ = probability of an effect due to the time of measurement, *P*_Int_ = probability of an interactive effect between R:B*_X_* blue content and measurement time.

**Figure 9.**
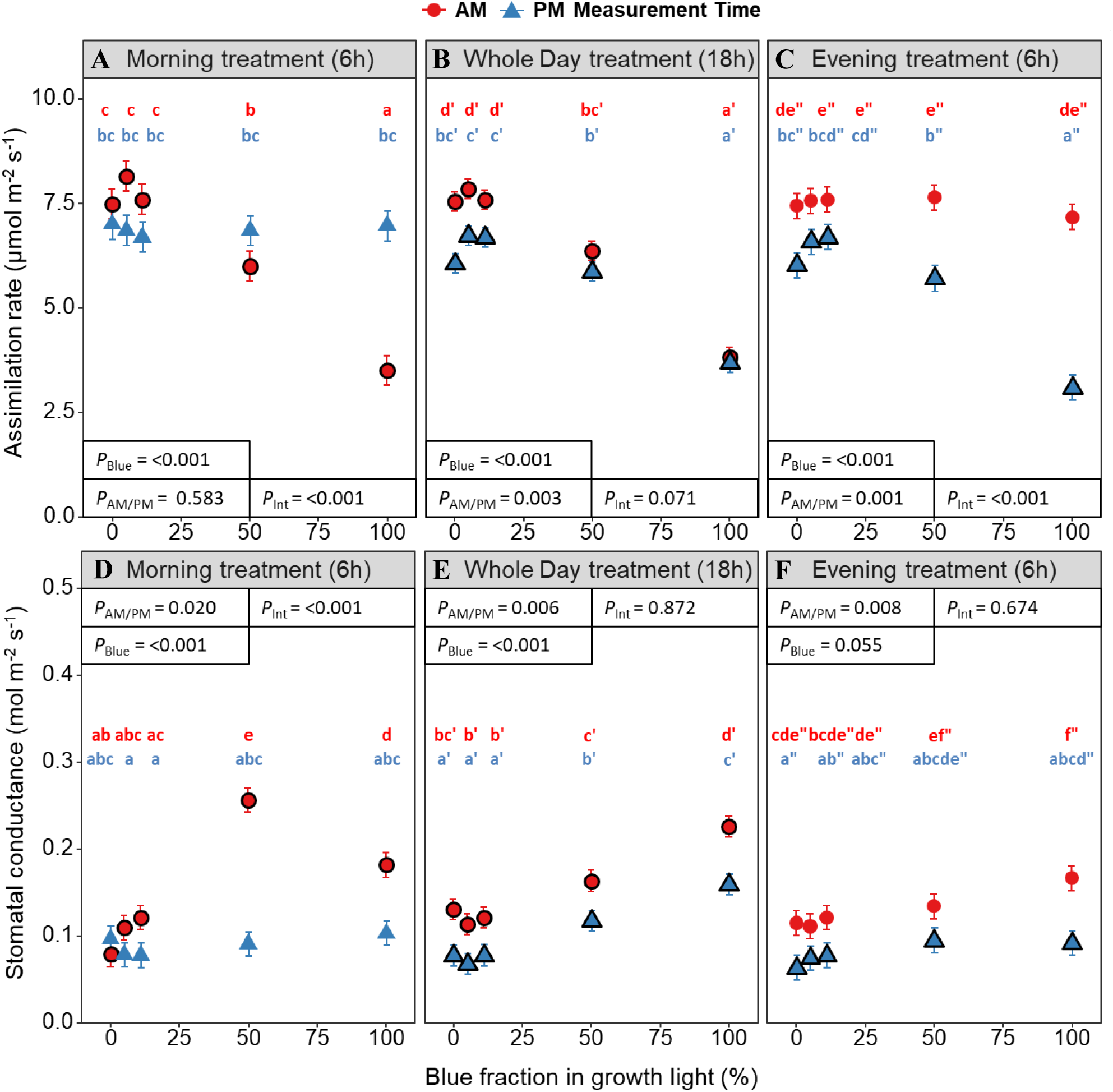
Assimilation rate and stomatal conductance of red lettuce grown under different red:blue treatments. Measurements during the Morning (AM) and Evening (PM) for assimilation rates (**A-C**; µmol CO_2_ m^−2^ s^−1^) and stomatal conductance (**D-F**; mol CO_2_ m^−2^ s^−1^) of lettuce cv. “Redflash” grown for 21 days. Five different R:B*_X_* ratios (R:B_100:0_, R:B_95:5_, R:B_89:11_, R:B_50:50_, and R:B_0:100_) were applied either during the six Morning hours, six Evening hours, or during Whole Day for all 18 hours of the day. Shape outlines indicate plants receiving the specific R:B*_X_* treatment during measurements, otherwise plants received R:B_89:11_. Datapoints represent treatment means and error bars represent standard error means of four growth cycles (*n* = 4), each with three replicate plants. Different letters indicate significantly different values for each combination of R:B*_X_* and measurement time, according to an unprotected Fisher LSD test (*a* = 0.05). Apostrophes indicate significance per treatment, for Morning (no apostrophe), Whole Day (′), and Evening (″). *P*_Blue_ = probability of an effect due to R:B*_X_* blue content, *P*_AM/PM_ = probability of an effect due to the time of measurement, *P*_Int_ = probability of an interactive effect between R:B*_X_* blue content and measurement time.

For Greenflash, each treatment period had a unique leaf assimilation pattern: 1) Morning treatments (**Figure 8A**), had reduced assimilation for R:B_0:100_ during AM measurements (while receiving R:B_0:100_), but this increased while receiving R:B_89:11_ during PM measurements; 2) Whole Day assimilation (**Figure 8B**) peaked for R:B_50:50_ plants during AM measurements and R:B_50:50_ and R:B_0:100_ during PM; and 3) Evening treatments (**Figure 8C**) increased assimilation with B fraction during AM, and during PM only R:B_50:50_ increased. Greenflash assimilation (**Figure 8A-C**) had interactive effects of B fraction and AM/PM measurement time for each of Morning (*p* = <0.001, *α* = 0.05), Whole Day (*p* = 0.009, *α* = 0.05), and Evening (*p* = 0.034, *α* = 0.05). The stomatal conductance was highest for all treatment periods during AM for both R:B_50:50_ and R:B_0:100_, which were always equal within treatment periods (**Figure 8A-C**). During PM measurements, conductance instead increased between R:B_50:50_ and R:B_0:100_ for Whole Day and Evening treatments. Stomatal conductance only had significant effects of B fraction and AM/PM measurement times for Whole Day and Evening treatments, not interactive effects (**Figure 8D-F**). Greenflash stomatal density increased with increased B fraction for Whole Day adaxial (*p* = 0.004), Whole Day abaxial (*p* = <0.001), and Morning abaxial leaf sides (*p* = <0.001; Table 1).

**Table 1.** Stomatal densities of adaxial and abaxial leaf sides of two lettuce cultivars grown under different R:B treatments.

| Leaf side | Period | R:B<br>100:0 | R:B<br>95:5 | R:B<br>89:11 | R:B<br>50:50 | R:B<br>0:100 | LSD | *P*<sub>Blue</sub> |
| --- | --- | --- | --- | --- | --- | --- | --- | --- |
| cv. "Greenflash" |  |  |  |  |  |  |  |  |
| Adaxial | Morning | 59<sup>a</sup> | 58<sup>a</sup> | 63<sup>a</sup> | 63<sup>a</sup> | 65<sup>a</sup> | 12.12 | 0.708 |
|  | Whole Day | 57<sup>b</sup> | 51<sup>b</sup> | 63<sup>ab</sup> | 78<sup>a</sup> | 73<sup>a</sup> | 15.04 | 0.004 |
|  | Evening | 74<sup>a</sup> | 59<sup>b</sup> | 63<sup>ab</sup> | 71<sup>ab</sup> | 74<sup>a</sup> | 13.35 | 0.124 |
| Abaxial | Morning | 51<sup>c</sup> | 60<sup>bc</sup> | 55<sup>c</sup> | 74<sup>a</sup> | 68<sup>ab</sup> | 8.63 | <0.001 |
|  | Whole Day | 56<sup>b</sup> | 50<sup>b</sup> | 55<sup>b</sup> | 71<sup>a</sup> | 81<sup>a</sup> | 10.03 | <0.001 |
|  | Evening | 60<sup>ab</sup> | 59<sup>ab</sup> | 55<sup>b</sup> | 71<sup>a</sup> | 68<sup>ab</sup> | 10.23 | 0.082 |
| cv. "Redflash" |  |  |  |  |  |  |  |  |
| Adaxial | Morning | 43<sup>b</sup> | 50<sup>b</sup> | 52<sup>b</sup> | 51<sup>b</sup> | 62<sup>a</sup> | 8.19 | 0.005 |
|  | Whole Day | 55<sup>bc3</sup> | 49<sup>c</sup> | 52<sup>bc</sup> | 62<sup>ab</sup> | 70<sup>a</sup> | 11.64 | 0.006 |
|  | Evening | 53<sup>ab</sup> | 49<sup>b</sup> | 52<sup>ab</sup> | 48<sup>b</sup> | 59<sup>a</sup> | 10.06 | 0.101 |
| Abaxial | Morning | 51<sup>c</sup> | 55<sup>bc</sup> | 62<sup>abc</sup> | 63<sup>ab</sup> | 72<sup>a</sup> | 11.77 | 0.011 |
|  | Whole Day | 55<sup>c</sup> | 53<sup>c</sup> | 62<sup>c</sup> | 81<sup>b</sup> | 97<sup>a</sup> | 12.63 | <0.001 |
|  | Evening | 78<sup>a</sup> | 80<sup>a</sup> | 62<sup>b</sup> | 72<sup>ab</sup> | 86<sup>a</sup> | 12.93 | 0.014 |

Note: Five different R:B<sub>x</sub> ratios (R:B<sub>100:0</sub>, R:B<sub>95:5</sub>, R:B<sub>89:11</sub>, R:B<sub>50:50</sub>, and R:B<sub>0:100</sub>) were applied either during the six Morning hours, six Evening hours, or during Whole Day for all 18 hours of the day. During the time no treatments were being applied, plants received a ratio of R:B<sub>89:11</sub>. Data are average values of four stomatal imprints from two growth cycles (*n* = 2). Data was tested for significance within leaf sides ( $\alpha$ = 0.05), so values are comparable within leaf sides. Means followed by different letters within each leaf side and treatment group differ significantly as determined by a Fisher least significance difference (LSD) test. *P*<sub>Blue</sub> = probability of an effect due to R:B<sub>x</sub> blue content.

For Redflash leaf assimilation, Morning treatments strongly decreased with increased B fraction, but only during AM measurements (while receiving R:B*_X_*)—during PM measurements (while receiving R:B_89:11_), there was no significant change. Whole Day assimilation (**Figure 9B**) showed the same strong decrease for both AM and PM measurements. Finally, Evening treatments (**Figure 9C**) had the opposite trend of Morning treatments, decreasing with increased B during PM measurements (while receiving R:B*_X_*), with no change during AM measurements (while receiving R:B_89:11_). Thus, both Morning and Evening had interactive effects of B fraction and AM/PM measurement time (both *p* = <0.001, *α* = 0.05). However, Whole Day had individual effects of B fraction and AM/PM measurement time (*p* = <0.001 and *p* = 0.003, *α* = 0.05), but no significant interaction (*p* = 0.071, *α* = 0.05). Like Greenflash, Redflash stomatal conductance of Morning, Whole Day, and Evening treatments all increased for both R:B_50:50_ and R:B_0:100_ during AM (**Figure 9D-F**). However, conductance did not significantly change during PM for Morning and Evening treatments, only increasing with B fraction for Whole Day treatments (**Figure 8B**). Of note, there was a strong peak in conductance for Morning treatments at R:B_50:50_, resulting in an interactive effect for Morning treatments (*p* = <0.001, *α* = 0.05). Whole Day and Evening conductance had no interactive effects, but Whole Day conductance increased with B fraction (*p* = <0.001, *α* = 0.05) and both Whole Day and Evening decreased under PM measurement times (*p* = 0.006 and *p* = 0.008, *α* = 0.05). Finally, Redflash stomatal density also increased with increasing B fraction for all treatments and leaf sides (Table 1), except for the adaxial stomata of the Evening treatment (*p* = 0.101; Table 1).

## 4 Discussion

### 4.1 Static monochromatic light leads to extreme morphology

In this study, we observed striking differences in morphological and photosynthetic traits when subjecting lettuce to both static R:B and diurnal spectral variation (different R:B ratios during the photoperiod). When grown under monochromatic R for the whole photoperiod, both Redflash and Greenflash showed long, curly leaves with extended petioles (**Figure 3**), traits attributed to the “red light syndrome” of plants subjected solely to monochromatic R light (Hogewoning et al., 2010; Miao et al., 2019; Trouwborst et al., 2016). Although these lettuce plants had greater tissue expansion than other treatments, their fresh weight, dry weight, and SLA did not differ from plants grown under R:B light with low B fraction (**Figure 3; Supplemental Table 1**). Furthermore, although plants with red light syndrome often have reduced photosynthetic capacity and stomatal function (Hogewoning et al., 2010), this study showed both parameters under R were the similar to plants grown in low B treatments (**Figure 8** and **Figure 9**). It was altogether unexpected that these plants grew considerably well (albeit with curly leaves and long petioles) and photosynthesized effectively under only R. Previous studies found much more negative impacts of monochromatic R, and although some have shown growth and fresh weight under monochromatic R can be similar to a combined R:B spectra, there were still apparent reductions in net photosynthesis (Tarakanov et al., 2022). To our awareness, this is the first study that shows less-affected photosynthesis in monochromatic R, potentially due to us measuring crops with instantaneous measurements with their treatment spectra, rather than measuring all plants with one R:B*_X_* spectrum. Monochromatic B also showed extreme traits previously seen in lettuce: decreased weight (Chen et al., 2021, 2019), fewer leaves (Chen et al., 2019; Saito et al., 2010), and hyponastic growth (Jishi et al., 2021a). Although this growth under monochromatic B is often simply stated to be elongated growth (Hernández and Kubota, 2016; Jishi et al., 2021b, 2021a), we suggest it is more specifically a hyponastic elongation in lettuce, as seen in *Arabidopsis* and tobacco (Keller et al., 2011; Pierik et al., 2004).

### 4.2 Diurnal variations in R:B ratio reduce the extreme morphology of static conditions

As R light contributes to various growth-promoting processes (Cammarisano et al., 2021; Kang et al., 2016; Wang et al., 2016) and B light is linked to enhanced nutritional content (Fasciolo et al., 2024; Samuolienė et al., 2017; Van Brenk et al., 2024), determining their combined effects is imperative to simultaneously improve plant growth and plant quality in VF. When monochromatic R or B was used for six hours with diurnal variation (irrespective of Morning or Evening), coupled with a high R:B for the remaining twelve hours of the photoperiod, the impacts of monochromatic R or B were less extreme (**Figure 3**). That is, neither the red-light syndrome, nor the hyponastic growth from monochromatic B, was seen for any plant grown under dynamic conditions (**Supplemental Figures 3 and 4**). Further, reducing the duration of high B in 18-hour static treatments to six hours—either in the morning or evening—caused plants to grow larger, with higher weight, leaf area, and leaf number (**Figure 2** and **Figure 3; Supplemental Figures 1-4; Supplemental Table 1**). This is of similar relevance to previous research that instead used a static combination of R and B to alleviate negative impacts of monochromatic R or B in cucumber (Miao et al., 2019). Here, we not only verified these findings with other static R and B combinations, but we also showed these monochromatic deficiencies can be mitigated through dynamic, diurnal treatments, as plants had twelve remaining photoperiod hours to compensate growth under low B and high R.

### 4.3 Assimilation and stomatal conductance are affected by instantaneous red:blue content

Leaf assimilation rate and stomatal conductance were higher in Greenflash than Redflash, which associated with the higher growth of Greenflash (**Figure 8** and **Figure 9**). Notably, compared to Greenflash, Redflash had a stronger decrease in assimilation and increase in conductance with high B; red lettuce leaf assimilation was consistently low with static high B exposure, agreeing with data from Wang et al. (2016), who also demonstrated that low R:B reduces assimilation in lettuce.

However, the most notable photosynthetic responses occurred in plants grown under diurnal variation. For both Morning and Evening treatments, leaf assimilation was low during periods of high B exposure but reverted to the level of standard conditions (R:B_89:11_) when being exposed to R:B_89:11_. This confirms the reversibility of responses to spectral changes found before in lettuce (Kim et al., 2004). Temporal differences were also observed, assimilation and stomatal conductance were both higher at the beginning of the photoperiod This aligns with numerous studies—including with lettuce—showing that photosynthesis is more active during the first part of a photoperiod (Horrer et al., 2016; Kaiser et al., 2019; Kim et al., 2004).

Increasing B fraction led to increased stomatal conductance (**Figure 8B** and **Figure 9B**), as shown previously (Kang et al., 2016; Kim et al., 2004; Pennisi et al., 2019), due to stomata having blue-light gated guard cells that open in response to blue light (Assmann et al., 1985; Kaiser et al., 2019).

Increased stomatal conductance may be due to stomatal density, which generally increased with increased B (Table 1). Higher stomatal density has been associated with greater stomatal conductance (Cano et al., 2019) and high B results in more stomata per cm^2^, as we show and others have shown that high B produces smaller leaves (Wang et al., 2016). Notably, we found that plants receiving high B during the evening had higher conductance than those under low B. Interestingly, the next morning, these plants grown under high B still had higher conductance, even under standard light (R:B_89:11_). Here, high B in the evening may actually promote stomatal opening for the next morning, potentially due to guard cell starch concentration. Briefly, in leaves under natural light conditions, starch accumulates during the photoperiod and is broken down overnight (Horrer et al., 2016; Zeeman et al., 2010). However, at the end of the night (beginning of the photoperiod), guard cells retain more starch than the rest of the leaf (Horrer et al., 2016), and this starch maintains stomatal closure. With the start of the photoperiod, B light causes a phototropin signalling cascade, activating enzymes that degrade guard cell starch, causing stomatal opening (Horrer et al., 2016). In our case, high B exposure during the evening may induce guard cell starch degradation prior to nighttime, which is further reduced overnight until morning, resulting in earlier opening of stomata and increased conductance.

### 4.4 Green lettuce grows larger under diurnal variations of high blue than with static high blue

Under diurnal variations, the total B fraction within a photoperiod was reduced compared to static conditions. When considering the fraction of total daily B received, Greenflash under 6 hours of monochromatic B had higher growth (fresh weight, dry weight, and leaf area) than what was projected by the trend of 18-hour static R:B*_X_* conditions (**Figure 2C; Supplemental Figures 1 and 2**). Conversely, Redflash grown under either dynamic conditions or static conditions followed the same decreasing growth trend with total daily B fraction (**Figure 2D**). This indicates that red lettuce growth appears to be wholly linked to the total B fraction received throughout the day, whereas green lettuce may more successfully grow under dynamic applications of high B. This may be due to the higher assimilation and stomatal conductance rates of Greenflash under high B, compared to Redflash. Under high B, Greenflash assimilation and stomatal conductance was less negatively affected than Redflash, resulting in similar weight and leaf area between Greenflash R:B_50:50_ and R:B_0:100_. On the other hand, the decreased assimilation of Redflash under high B resulted in decreased weight and leaf area between R:B_50:50_ to R:B_0:100_. Therefore, the comparatively greater growth of Greenflash under high B, when transitioned to low B for 12 hours in dynamic conditions, may have had a greater capacity to compensate, expand, and grow further during these remaining hours of the photoperiod.

### 4.5 Chlorophyll and carotenoid content is reduced with more than 50% total daily blue fraction

Some plant pigments such as chlorophyll and carotenoids have been found to scavenge ROS, and carotenoids can also provide photoprotection (Maoka, 2020; Zulfiqar et al., 2021). We initially postulated that darker green leaves under high B may have been due to increased chlorophyll or carotenoids, as these are pigments that produce green or yellow colouration and increase under B exposure (Samuolienė et al., 2017; Van Brenk et al., 2024). In the present study, chlorophyll was highest at 50% B, and carotenoid content decreased with high B (≥50%), for both static and (to a lesser extent) dynamic light conditions. To note, prior studies used total B fractions below 50%. Thus, there is likely a fraction of B light—above 11% and below 50%—that maximizes chlorophyll and carotenoid production, in line with suggestions from Samuolienė et al. (2017). Finally, certain carotenoids such as zeaxanthin and lutein make leaves yellower (Khoo et al., 2011), so low B may cause leaves to appear lighter in colour. This, in conjunction with a greater relative content of green chlorophyll over yellow carotenoids, can cause leaves grown with high B to appear darker green.

### 4.6 Red lettuce anthocyanins and flavonoids are produced more effectively with high blue fraction in diurnal variation

The changes in pigmentation of both cultivars are likely strongly linked to their identifiable differences in the metabolomic profiles of high B conditions compared with low B conditions (**Figure 4**). In red lettuce, this red pigmentation is attributed to anthocyanin accumulation, red-purple pigments which have upregulated production during periods of stress (Dixon and Paiva, 1995; Sarkar and Shetty, 2014; Van Brenk et al., 2024). Anthocyanins and other flavonoids are a subclass of phenylpropanoid compounds associated with photoprotection and ROS scavenging (Dixon and Paiva, 1995; Falcone Ferreyra et al., 2012; Panche et al., 2016; Williams et al., 2004). In this study, anthocyanins likely accumulated in response to the high-energy B light, as they are high-energy light-filtering antioxidants. Most remarkably, anthocyanins and flavonoids seemed to respond to the highest B fraction of instantaneous light received rather than by the average R:B ratio of a day, independent of the timing of diurnal application. That is, six hours of diurnal treatments with high B produced roughly the same anthocyanin content as 18 hours of high B. Flavonoids showed similar results, as they have also been found to increase with high B (Sarkar & Shetty, 2014). It is incredibly valuable to producers that anthocyanin and flavonoid content with diurnal spectral variations exceeded the trend of static treatments. This means that high B exposure to induce the production of these nutritionally relevant compounds can be dramatically reduced to one-third of the photoperiod, leaving the remaining two-thirds of the off-treatment photoperiod to be used to benefit growth.

### 4.7 Green lettuce flavonoids are readily produced under high blue light

The change in leaf pigmentation from light to dark green under increased B for Greenflash is not due to anthocyanins, as this green lettuce does not produce anthocyanins (**Supplemental Figure 5**).

However, there was still a clear separation between high B exposure and low B exposure when considering the metabolic profiles of green lettuce (**Figure 4**). We postulate that the B-induced changed pigmentation in Greenflash may be linked to the strong increase of total flavonoid content with increased B (likely also applicable to Redflash, to some extent). While both green and red lettuce increased flavonoid content with B fraction, green lettuce had a notably steeper increase, to a plateau. Therefore, green lettuce may more readily produce more antioxidant flavonoids to an optimum, in order to combat light-induced stress through their antioxidant capacity (Falcone Ferreyra et al., 2012; Shi et al., 2022). This is further substantiated as Greenflash produced more overall flavonoids than Redflash, potentially accommodating for its lack of anthocyanins. It is possible that some colour-producing flavonoids—such as the yellow colour-causing quercetin (Anand David et al., 2016)—may accumulate in greater quantities in green lettuce via phenylpropanoid biosynthesis pathways that can branch to produce anthocyanins (in red lettuce) but are unable to go further than their flavonoid precursors in green lettuce (Wada et al., 2022).

### 4.8 Growth and antioxidant production trade-offs may be based on inherent plant pigmentation

When considering total daily B fraction, Greenflash growth under diurnal high B exceeded the trends of static R:B treatments, whereas Redflash flavonoid and anthocyanin concentrations under diurnal high B exceeded the trend of static R:B. These results present an interesting consideration for how lettuce responds to high B—green lettuce and red lettuce differentially prioritize their responses to dynamic applications of high B light. Although both are negatively impacted by high B, they show different modes of recovery when transitioned to a low-stress low B environment. Green lettuce, after a transition from high B to low B, prioritizes growth and expands more while it has the opportunity to do so. Conversely, red lettuce rather prioritizes achieving protective needs, allowing for healthier photosystems, at the cost of the growth. This is further supported as Greenflash had consistently higher assimilation than Redflash, especially under the high B conditions wherein Redflash had strongly reduced assimilation. These negative repercussions on red lettuce assimilation under high B are likely due to a cycle of energy-prioritization that is tightly linked with pigment production. In response to high B, red lettuce produces more anthocyanins to filter high-energy light, protecting itself. The increased anthocyanin production results in a more concentrated mesh of these pigments, leading to more light-filtering. Thus, less light energy can be received and utilized by the photosynthetic apparatus, reducing the energy and carbon harnessed from photosynthesis. Then, the limited energy and carbon that *is* made accessible through photosynthesis is further divided again, being allocated to simultaneously produce more photoprotective pigments and growth-related compounds. This results in a positive feedback loop, promoting pigment production at the expense of growth for red lettuce.

## 5 Conclusions

The field of light spectra research in vertical farms continues to advance, offering new opportunities to optimize the growth of many crops in controlled environments. Here, we report that although high B reduces plant growth compared to low B, diurnal high B exposure improved plant growth over static conditions, which we attribute to instantaneous R:B light effects on leaf photosynthesis. High B reduced photosynthesis only during the six hours of high B exposure, then photosynthesis was at a regular level for the remaining 12 hours under low B. This corresponded with our findings that growth is largely related to daily B fraction, rather than what time B is distributed within a day. In contrast, anthocyanins and flavonoids in red lettuce responded to the highest B fraction received, regardless of if that R:B ratio was applied for six hours diurnally in the morning or evening, or for 18 hours of the whole day. The application of these diurnal variations of R:B ratios was presented as a method to reduce the duration of static high B treatments.

## 6 Data Availability

The data underlying this article will be shared on reasonable request to the corresponding author.

## 8 Acknowledgments

We would like to thank the Unifarm Klima staff, namely Gerrit Stunnenberg, Chris van Asselt, David Brink, and Dieke Smit. We would also like to thank Yasmin Dijksterhuis, Fleur Gulien, and Sarah Berman for their assistance with the experiment and Elias Kaiser for his input for the experimental setup. Additionally, we would like to thank Ep Heuvelink for his assistance with statistical analysis.

## 9 Author Contributions

**Jordan Van Brenk:** Conceptualization, Methodology, Formal analysis, Investigation, Writing - Original Draft, Writing - Review & Editing, Visualization, Supervision. **Kimberly Vanderwolk:** Methodology, Formal analysis, Investigation, Writing - Review & Editing. **Sumin Seo:** Formal analysis, Investigation, Writing - Review & Editing, Visualization. **Young Hae Choi:** Methodology, Writing - Review & Editing, Supervision. **Leo Marcelis:** Writing - Review & Editing, Supervision, Project administration, Funding acquisition. **Julian Verdonk:** Methodology, Writing - Review & Editing, Supervision, Project administration.

