## Supplemental Data for "Blue Light Sonata: Dynamic variation of red:blue ratio during the photoperiod differentially affects leaf photosynthesis, pigments, and growth in lettuce"

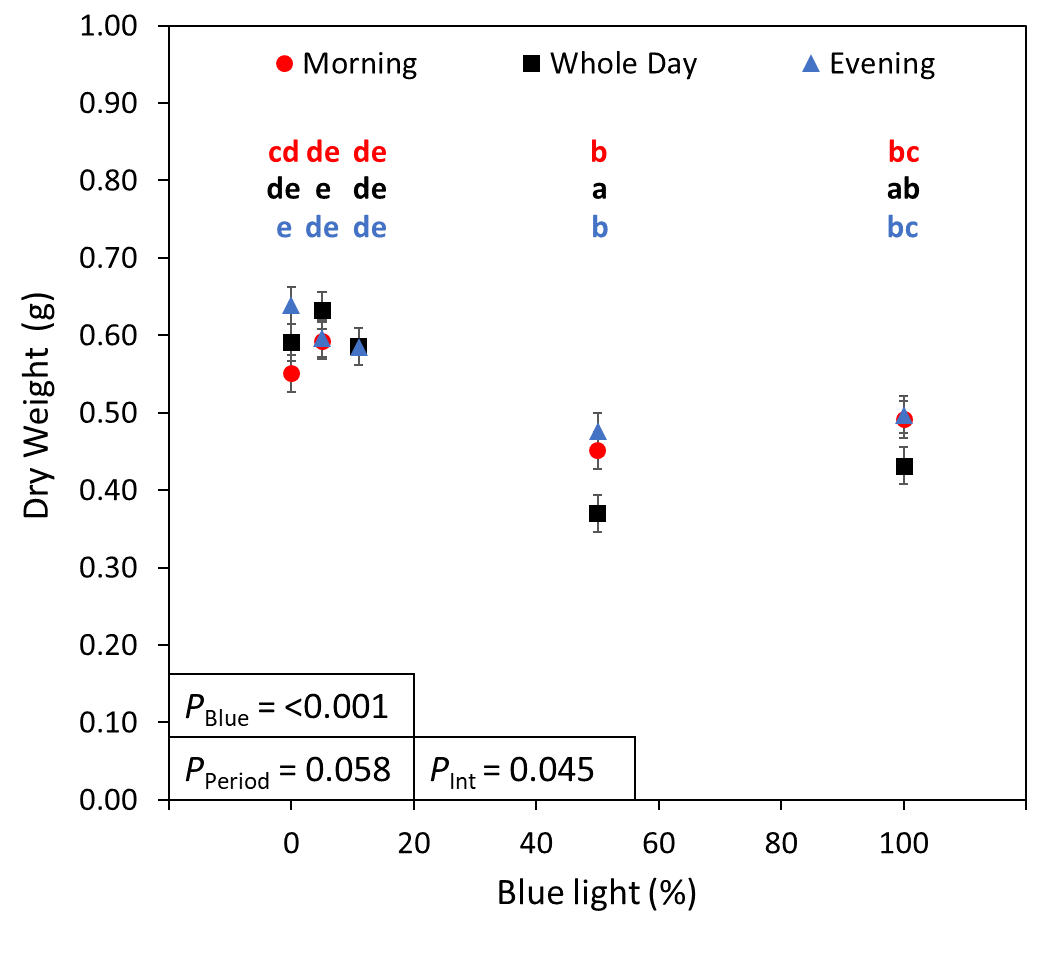

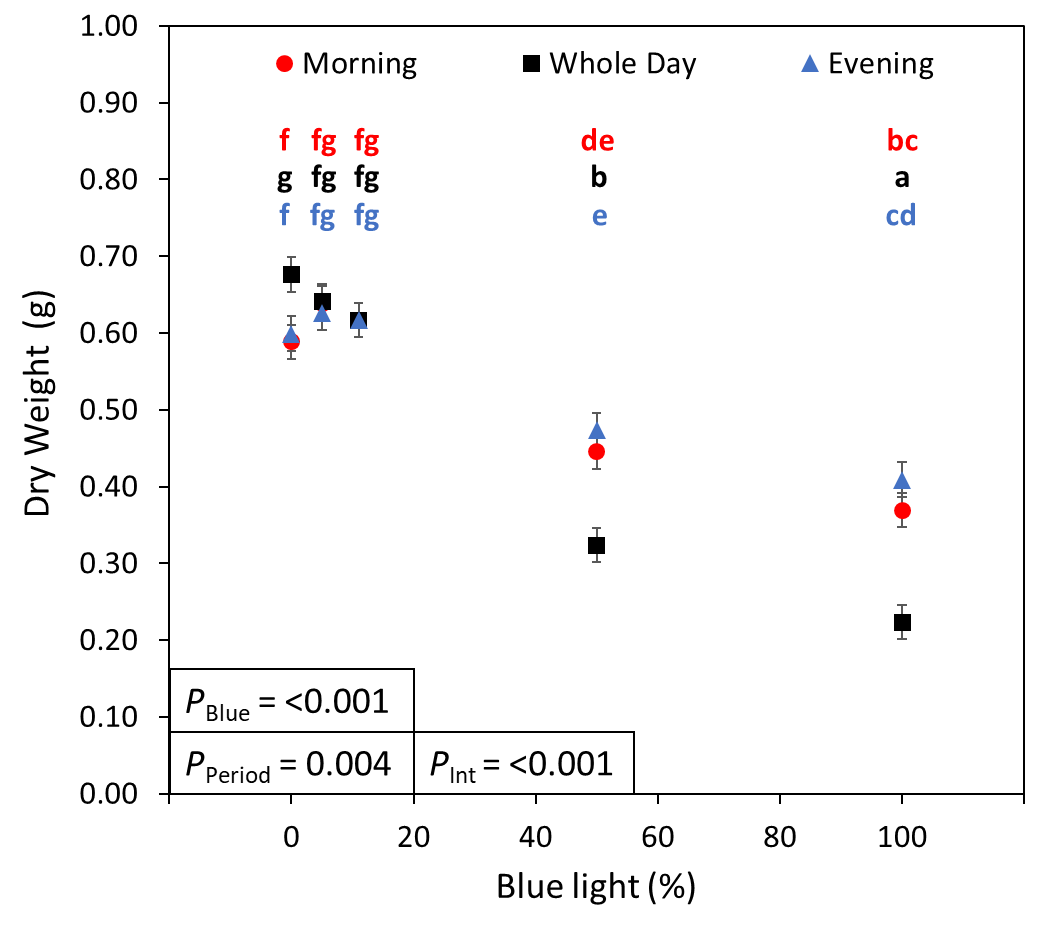

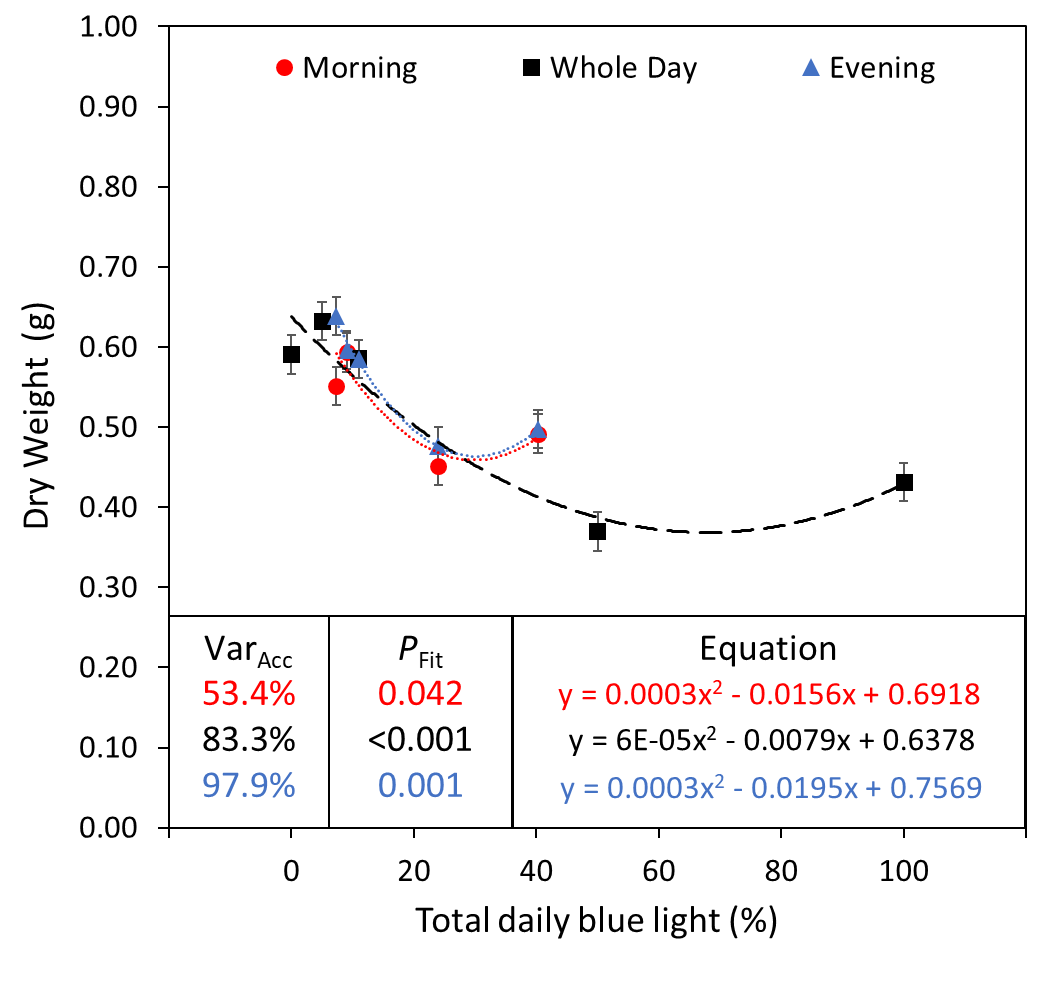

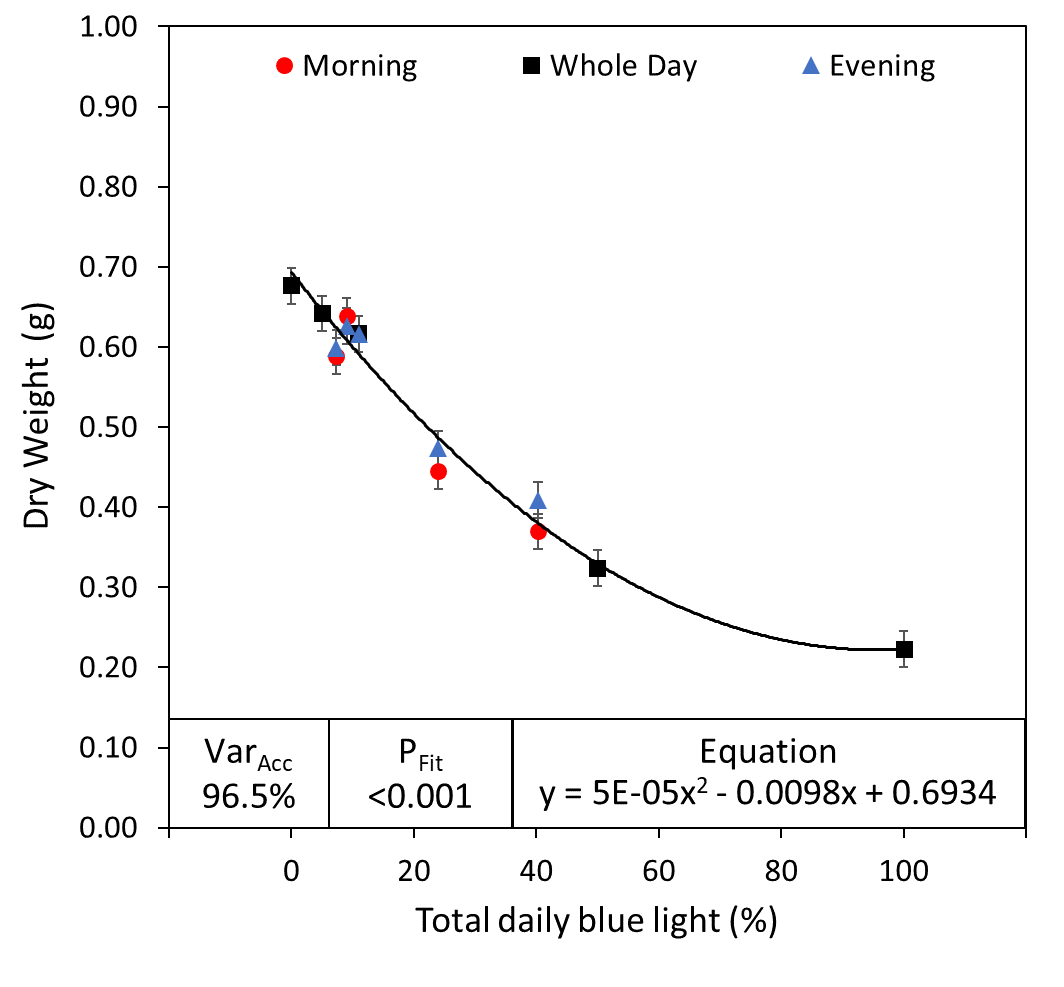

**C**

**D**

**B**

**A**

**cv. “Greenflash”**

**cv. “Redflash”**

Supplemental Figure 1. Dry weight of two lettuce cultivars grown under five different red:blue ratios applied during three different parts of the day.

Dry weight (g) of lettuce cv. “Greenflash” (**A, C**) and cv. “Redflash” (**B, D**) grown for 21 days. Five different R:B*_X_* ratios (R:B_100:0_, R:B_95:5_, R:B_89:11_, R:B_50:50_, and R:B_0:100_) were applied either during the six Morning hours, six Evening hours, or during Whole Day for all 18 hours of the day. The fraction of B during the treatment period (**A, B**) or total B fraction for the whole photoperiod (**C, D**) were used as X-axes. Datapoints represent means with standard error means of four growth cycles (*n* = 4), each consisting of sixteen replicate plants. For panels (**A**) and (**B**), different letters indicate significantly different values for each combination of R:B*_X_* and treatment period, according to an unprotected Fisher LSD Test (*a* = 0.05). *P*_Blue_ = probability of an effect due to R:B*_X_* blue content; *P*_Period_ = probability of an effect due to the treatment period of Morning, Whole Day, or Evening; *P*_int_ = probability of an interactive effect between R:B*_X_* blue content and treatment period; Var_Acc_ = percent variance accounted for by regression line; *P*_Fit_ = probability of a linear or quadratic trend.

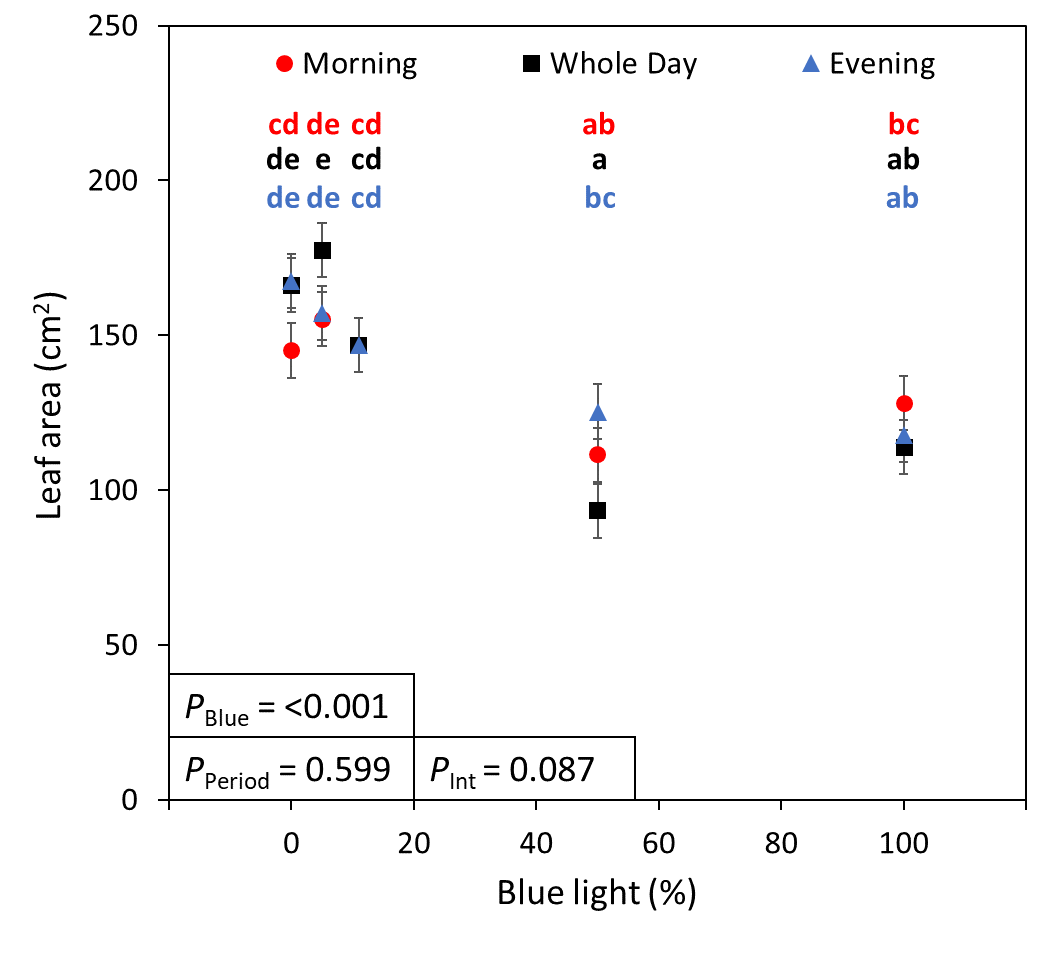

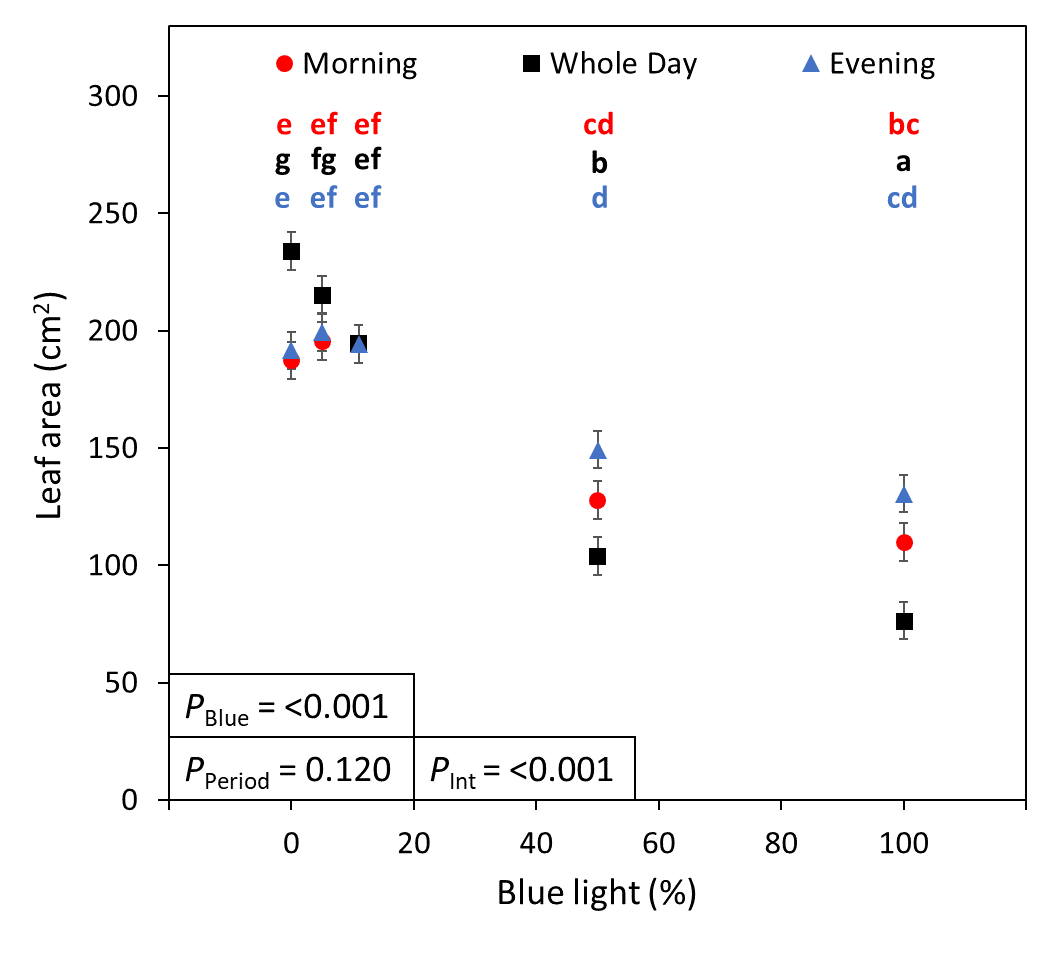
s
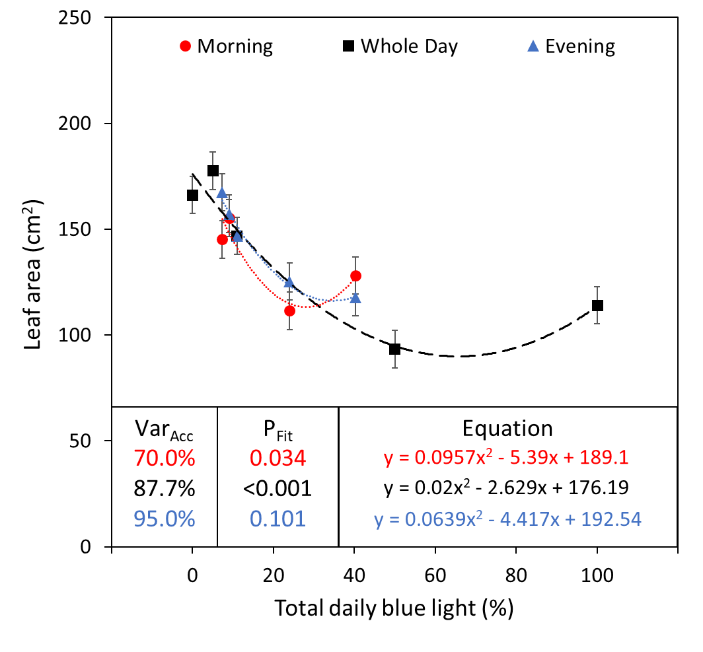

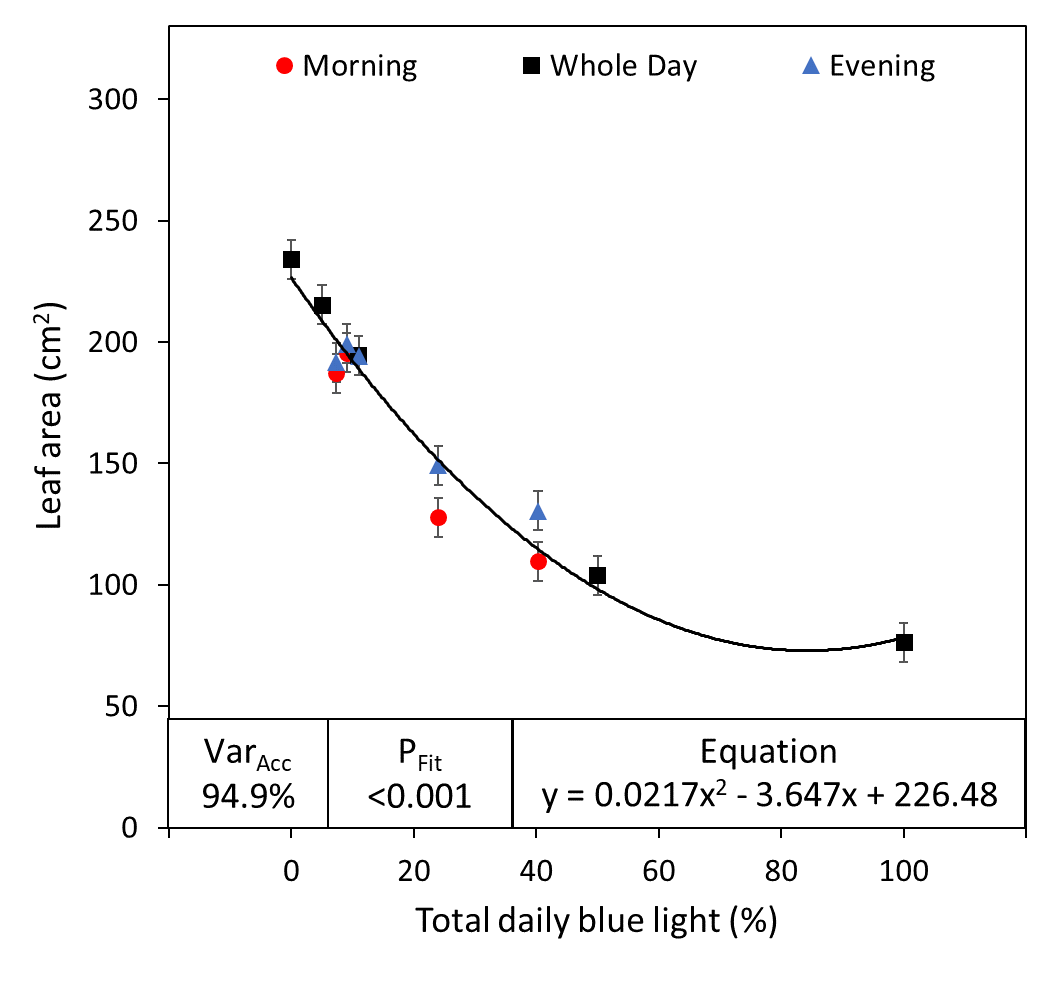

**B**

**D**

**cv. “Greenflash”**

**cv. “Redflash”**

**C**

**A**

Supplemental Figure 2. Leaf area of two lettuce cultivars grown under five different red:blue ratios applied during three different parts of the day.

Leaf area (cm^2^) of lettuce cv. “Greenflash” (**A, C**) and cv. “Redflash” (**B, D**) grown for 21 days. Five different R:B*_X_* ratios (R:B_100:0_, R:B_95:5_, R:B_89:11_, R:B_50:50_, and R:B_0:100_) were applied either during the six Morning hours, six Evening hours, or during Whole Day for all 18 hours of the day. The fraction of B during the treatment period (**A, B**) or total B fraction for the whole photoperiod (**C, D**) were used as X-axes. Datapoints represent means with standard error means of four growth cycles (*n* = 4), each consisting of sixteen replicate plants. For panels (**A**) and (**B**), different letters indicate significantly different values for each combination of R:B*_X_* and treatment period, according to an unprotected Fisher LSD Test (*a* = 0.05). *P*_Blue_ = probability of an effect due to R:B*_X_* blue content; *P*_Period_ = probability of an effect due to the treatment period of Morning, Whole Day, or Evening; *P*_int_ = probability of an interactive effect between R:B*_X_* blue content and treatment period; Var_Acc_ = percent variance accounted for by regression line; *P*_Fit_ = probability of a linear or quadratic trend.

Supplemental Table 1. Leaf traits and pigments in lettuce grown under different light spectra.

| Parameter | **Period** |  | **R:B**  **^100:0^** | **R:B**  **^95:5^** | **R:B**  **^89:11^** | **R:B**  **^50:50^** | **R:B**  **^0:100^** | **SEM^ǂ^** | ***P*_Blue_^ǂǂ^** | ***P*_Period_^ǂǂǂ^** | ***P*_Int_** |
| --- | --- | --- | --- | --- | --- | --- | --- | --- | --- | --- | --- |
| cv. “Greenflash” | | | | | | | | | | | |
| No. of leaves (# plant^-1^) | Morning |  | 9.72^ef^ | 9.89^f^ | 9.42^def^ | 9.12^cd^ | 8.82^bc^ | ± 0.191 | <0.001^*^ | <0.001^*^ | <0.001^*^ |
|  | Whole Day |  | 9.64^def^ | 9.48^def^ | 9.42^def^ | 8.69^bc^ | 7.06^a^ |  |  |  |  |
|  | Evening |  | 9.48^def^ | 9.38^def^ | 9.42^def^ | 9.19^cde^ | 8.36^b^ |  |  |  |  |
| Specific leaf area (cm^2^ g^-1^) | Morning |  | 262.7^bcd^ | 256.5^ab^ | 252.8^ab^ | 249.7^ab^ | 260.4^bc^ | ± 6.71 | 0.027^*^ | 0.029^*^ | 0.059 |
|  | Whole Day |  | 278.7^cd^ | 280.5^d^ | 252.8^ab^ | 252.7^ab^ | 265.4^bcd^ |  |  |  |  |
|  | Evening |  | 259.3^b^ | 261.8^bcd^ | 252.8^ab^ | 265.1^bcd^ | 237.8^a^ |  |  |  |  |
| Chlorophyll a  (mg g^-1^ DW) | Morning |  | 5.99^bc^ | 5.95^bc^ | 6.28^cd^ | 6.34^cd^ | 5.74^ab^ | ± 0.147 | <0.001^*^ | 0.197 | 0.105 |
|  | Whole Day |  | 5.41^a^ | 5.71^ab^ | 6.28^cd^ | 6.55^d^ | 5.79^ab^ |  |  |  |  |
|  | Evening |  | 6.11^bc^ | 6.10^bc^ | 6.28^cd^ | 6.34^cd^ | 5.77^ab^ |  |  |  |  |
| Chlorophyll b  (mg g^-1^ DW) | Morning |  | 3.25^bcd^ | 3.27^bcd^ | 3.43^de^ | 3.54^ef^ | 3.16^b^ | ± 0.072 | <0.001^*^ | <0.001^*^ | <0.001^*^ |
|  | Whole Day |  | 2.90^a^ | 3.09^ab^ | 3.43^de^ | 3.73^f^ | 3.28^bcd^ |  |  |  |  |
|  | Evening |  | 3.41^de^ | 3.39^cde^ | 3.43^de^ | 3.55^ef^ | 3.20^bc^ |  |  |  |  |
| $\frac{Chl a}{Chl b}$ | Morning |  | 1.839^defg^ | 1.816^cdef^ | 1.831^defg^ | 1.785^abc^ | 1.818^cdef^ | ± 0.0144 | <0.001^*^ | 0.243 | 0.006^*^ |
|  | Whole Day |  | 1.865^g^ | 1.847^fg^ | 1.831^defg^ | 1.754^a^ | 1.765^ab^ |  |  |  |  |
|  | Evening |  | 1.792^abcd^ | 1.800^bcde^ | 1.831^defg^ | 1.785^abc^ | 1.802^bcde^ |  |  |  |  |
| cv. “Redflash” | | | | | | | | | | | |
| No. of leaves (# plant^-1^) | Morning |  | 7.13^ef^ | 7.12^ef^ | 7.09^ef^ | 6.93^de^ | 6.57^bc^ | ± 0.100 | <0.001^*^ | <0.001^*^ | <0.001^*^ |
|  | Whole Day |  | 6.95^def^ | 6.89^de^ | 7.09^ef^ | 6.41^b^ | 5.09^a^ |  |  |  |  |
|  | Evening |  | 7.09^ef^ | 7.22^f^ | 7.09^ef^ | 6.80^cd^ | 6.38^b^ |  |  |  |  |
| Specific leaf area (cm^2^ g^-1^) | Morning |  | 318.7^bcd^ | 307.7^ab^ | 318.2^bcd^ | 287.7^a^ | 299.6^ab^ | ± 8.62 | 0.065 | <0.001^*^ | 0.325 |
|  | Whole Day |  | 349.4^e^ | 337.5^cde^ | 318.2^bcd^ | 322.3^bcd^ | 342.2^de^ |  |  |  |  |
|  | Evening |  | 321.3^bcd^ | 320.4^bcd^ | 318.2^bcd^ | 315.4^bc^ | 322.0^bcd^ |  |  |  |  |
| Chlorophyll a  (mg g^-1^ DW) | Morning |  | 3.13^ab^ | 3.31^bcd^ | 3.22^abcd^ | 3.26^bcd^ | 3.05^a^ | ± 0.075 | 0.017^*^ | 0.383 | 0.094 |
|  | Whole Day |  | 3.08^a^ | 3.15^abc^ | 3.22^abcd^ | 3.41^d^ | 3.41^d^ |  |  |  |  |
|  | Evening |  | 3.14^ab^ | 3.22^abcd^ | 3.22^abcd^ | 3.36^cd^ | 3.31^bcd^ |  |  |  |  |
| Chlorophyll b  (mg g^-1^ DW) | Morning |  | 1.53^abc^ | 1.64^def^ | 1.58^bcd^ | 1.69^efg^ | 1.60^bcde^ | ± 0.035 | <0.001^*^ | 0.013^*^ | <0.001^*^ |
|  | Whole Day |  | 1.45^a^ | 1.51^ab^ | 1.58^bcd^ | 1.87^h^ | 1.98^i^ |  |  |  |  |
|  | Evening |  | 1.58^bcd^ | 1.62^cdef^ | 1.58^bcd^ | 1.72^fg^ | 1.76^g^ |  |  |  |  |
| $\frac{Chl a}{Chl b}$ | Morning |  | 2.041^fg^ | 2.017^efg^ | 2.039^fg^ | 1.934^cd^ | 1.906^c^ | ± 0.0268 | <0.001^*^ | 0.288 | <0.001^*^ |
|  | Whole Day |  | 2.129^h^ | 2.081^gh^ | 2.039^fg^ | 1.828^b^ | 1.716^a^ |  |  |  |  |
|  | Evening |  | 1.991^def^ | 1.983^def^ | 2.039^fg^ | 1.946^cde^ | 1.880^bc^ |  |  |  |  |

Note: The R:B values are the five R:B*_X_* treatment ratios used in this study. These ratios were applied either during the six Morning hours, six Evening hours, or during Whole Day for all 18 hours of the day. Different letters indicate significantly different values for each combination of R:B*_X_* and treatment period, according to an unprotected Fisher LSD Test (*a* = 0.05). *P*_Blue_ = probability of an effect due to R:B*_X_* blue content; *P*_Period_ = probability of an effect due to the treatment period of Morning, Whole Day, or Evening. *P*_int_ = probability of an interactive effect between R:B*_X_* blue content and treatment period.

R:B_100:0_

R:B_95:5_

R:B_89:11_

R:B_50:50_

R:B_0:100_

Supplemental Figure 3. Representative photos of green lettuce cv. “Greenflash” grown under one of five R:B*_X_* treatments for Morning, Whole Day, and Evening Treatments

Representative photos of lettuce cv. “Greenflash” grown under Morning (top), Whole Day (middle), and Evening (bottom) treatments of one of five R:B_X_ treatments: (left to right) R:B_100:0_, R:B_95:5_, R:B_89:11_, R:B_50:50_, R:B_0:100_ . For all panels: upper photos are side-view representative photos, lower photos are overhead representative photos. Whole Day and Evening R:B_89:11_ plants were grown together, hence the same treatment/photograph.

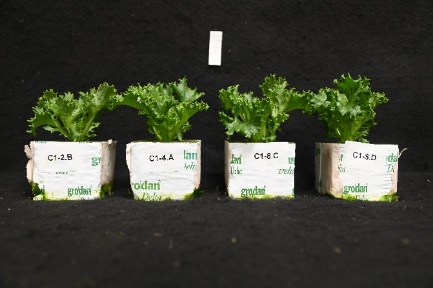

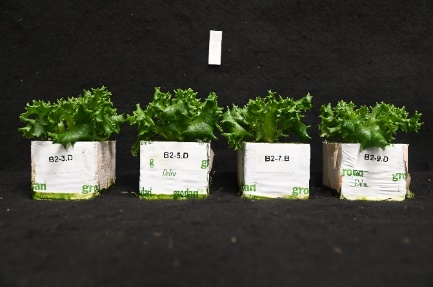

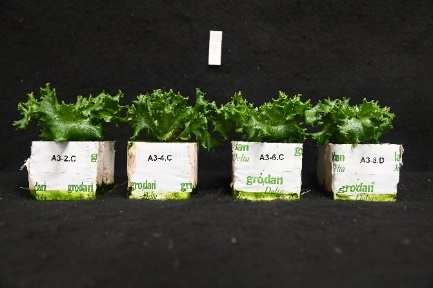

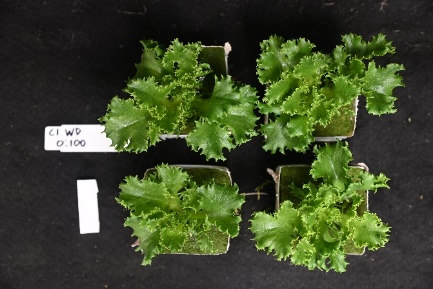

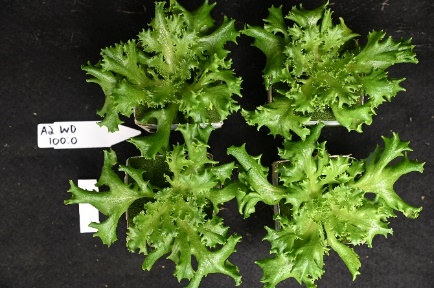

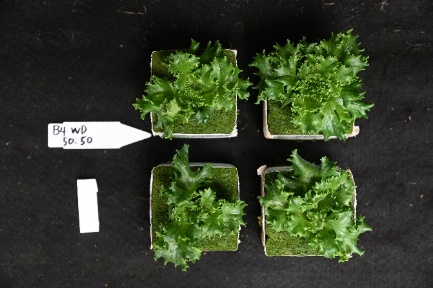

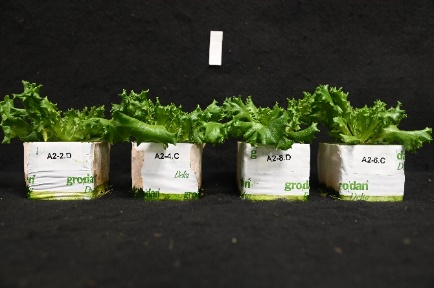

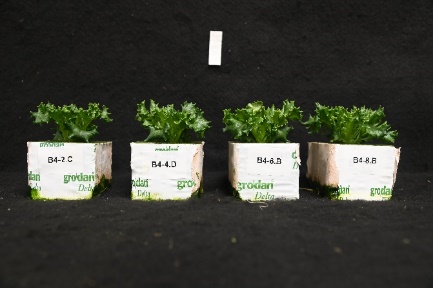

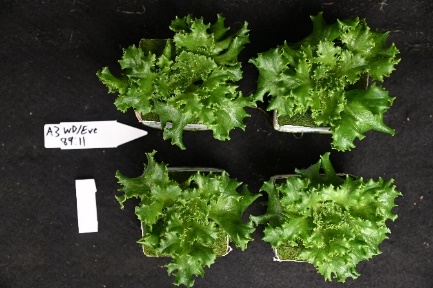

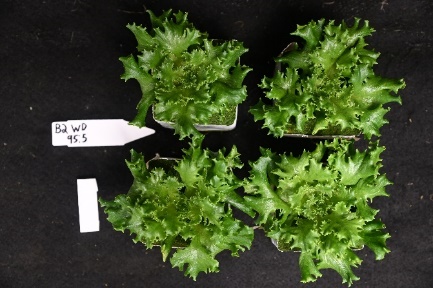

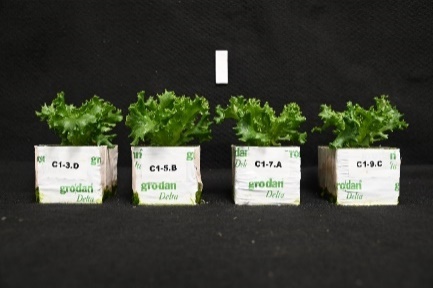

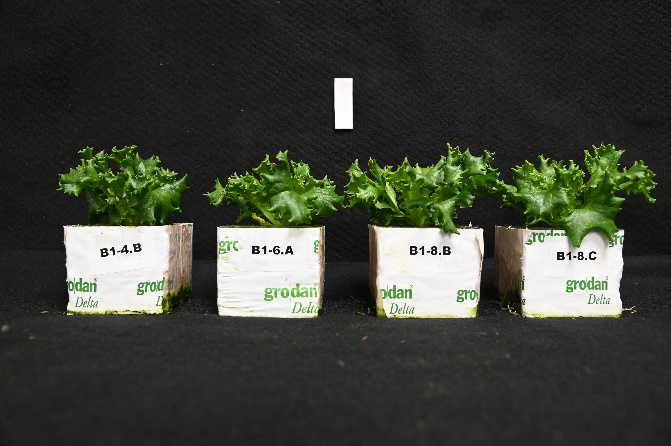

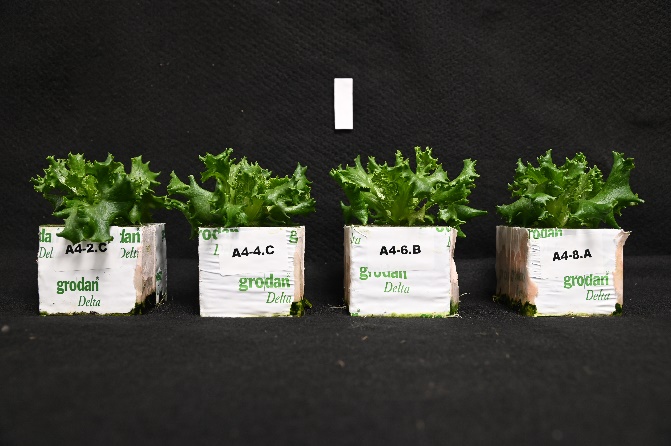

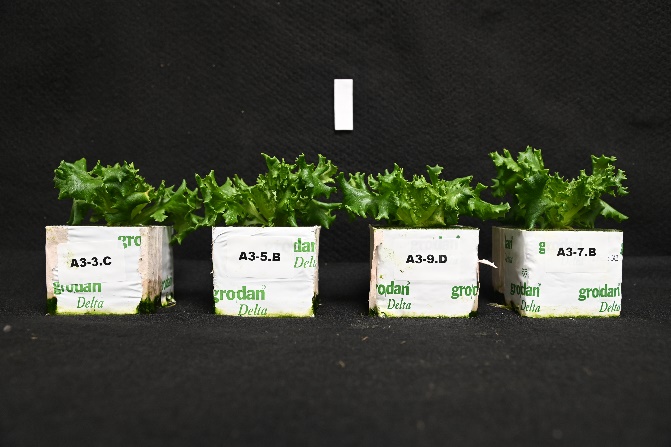

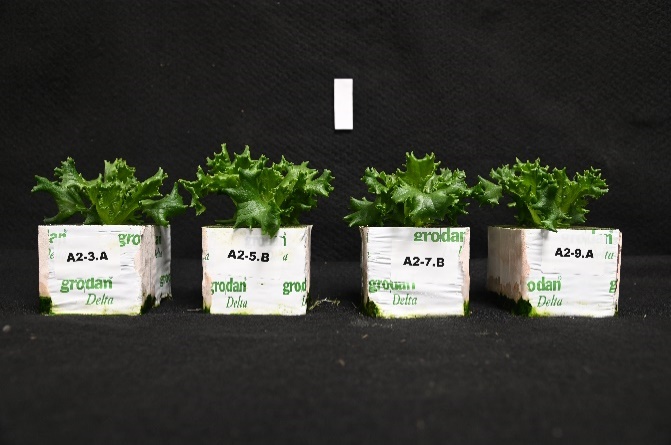

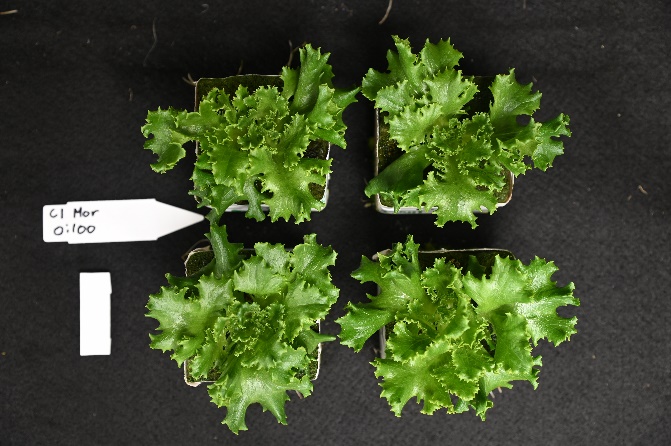

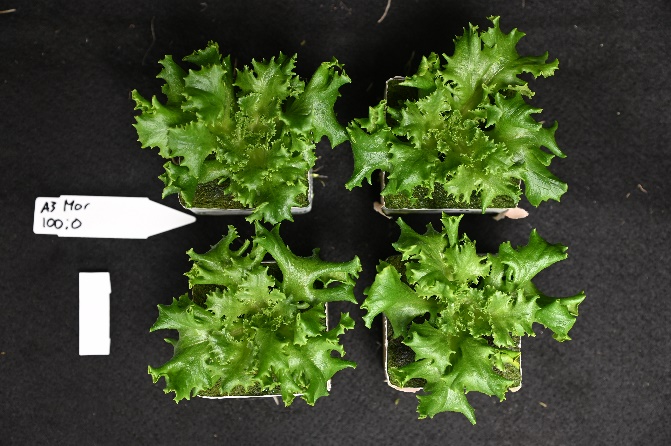

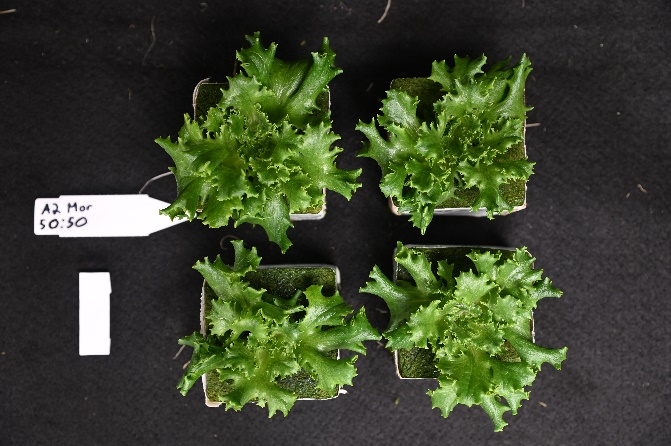

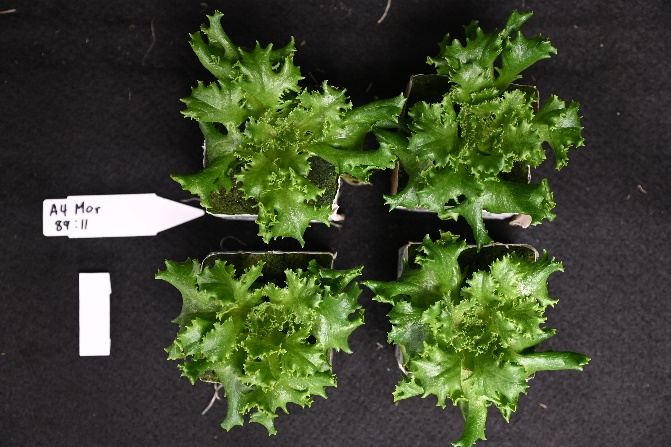

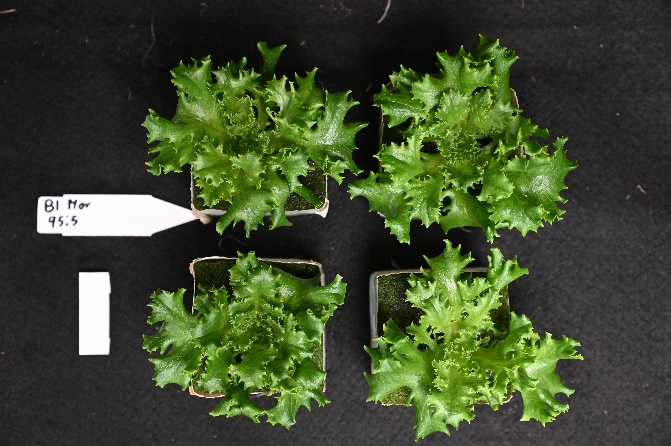

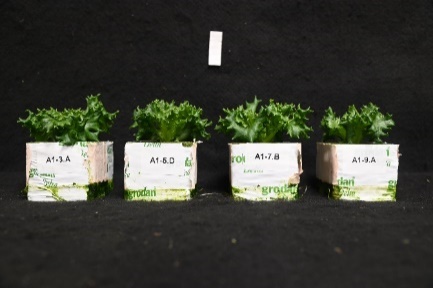

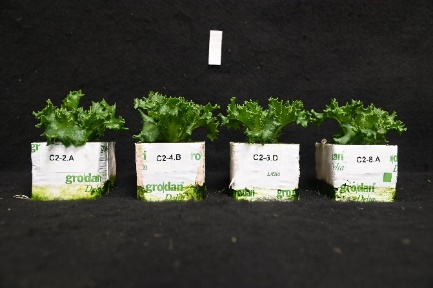

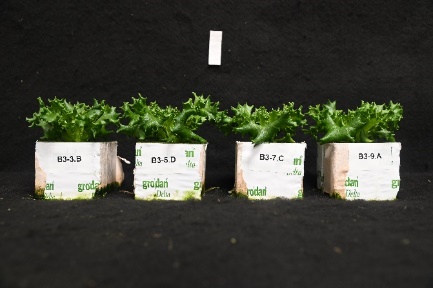

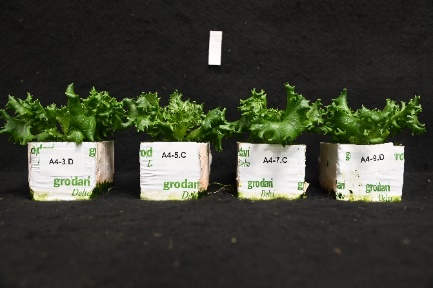

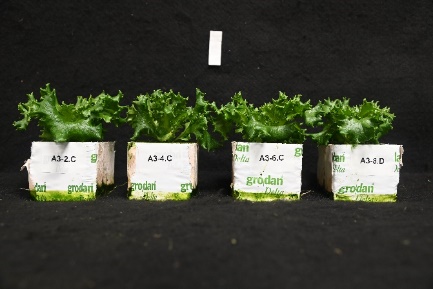

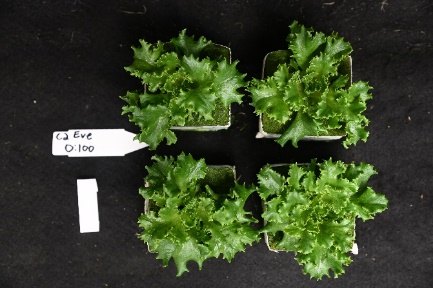

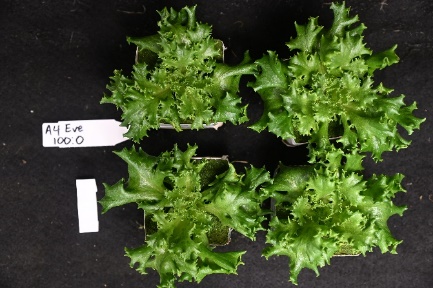

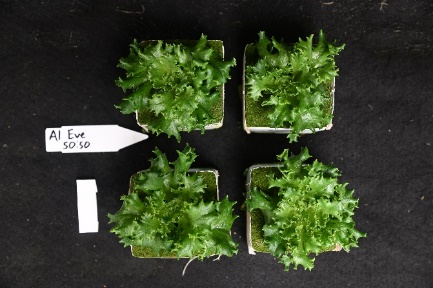

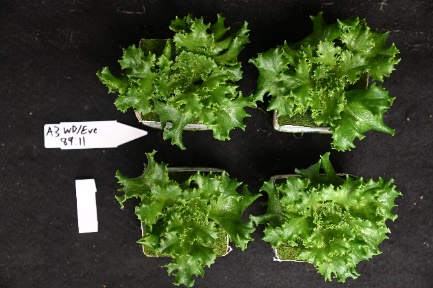

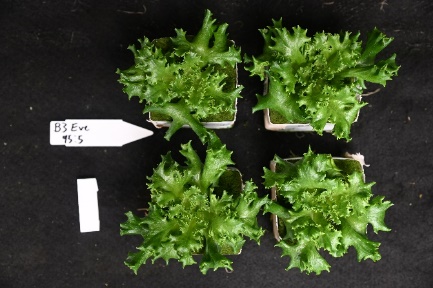

R:B_100:0_

R:B_95:5_

R:B_89:11_

R:B_50:50_

R:B_0:100_

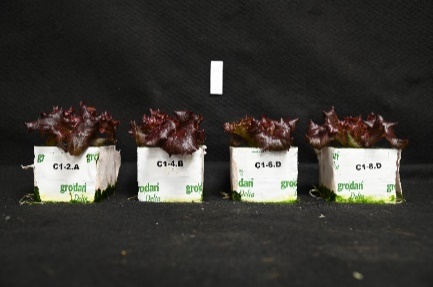

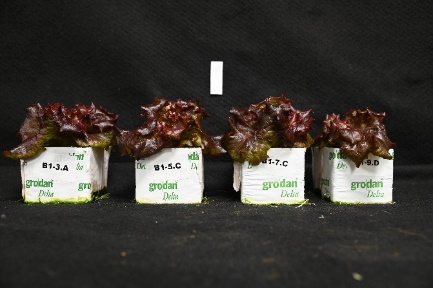

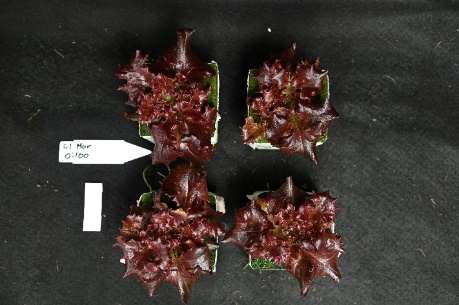

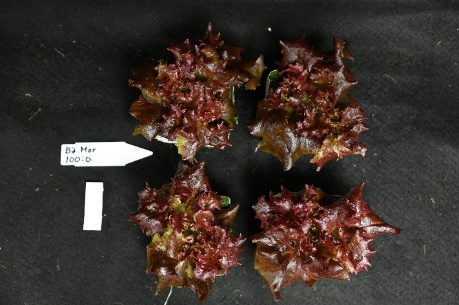

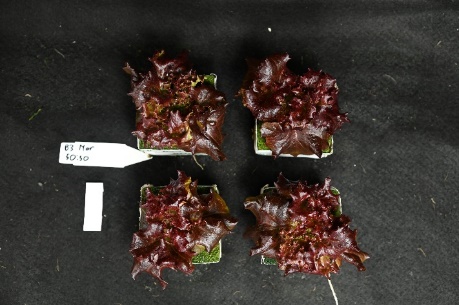

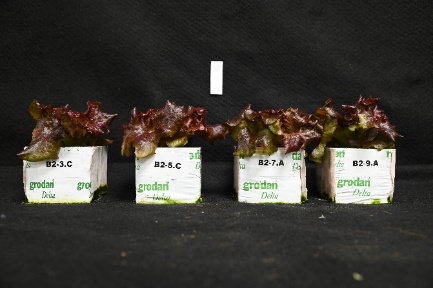

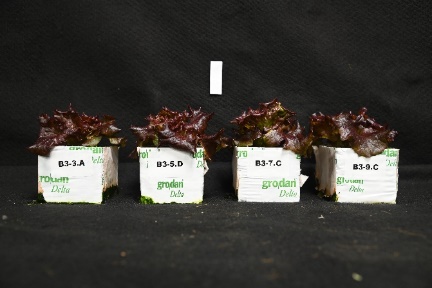

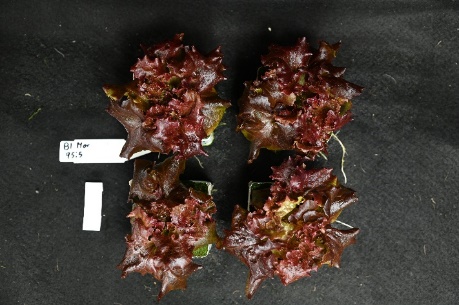

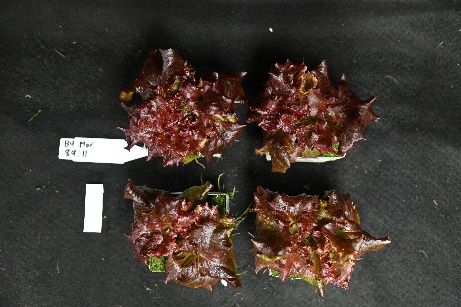

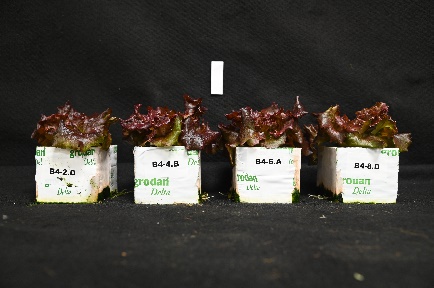

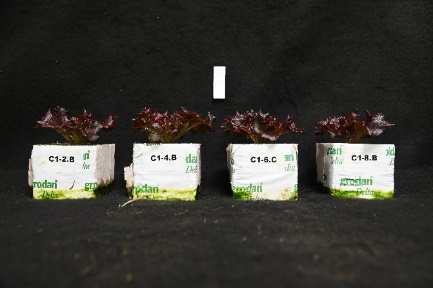

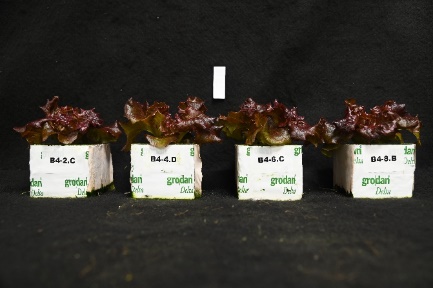

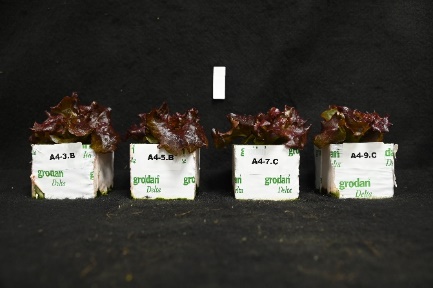

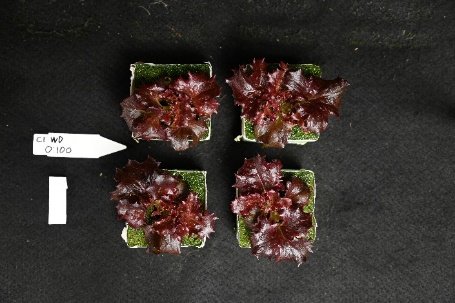

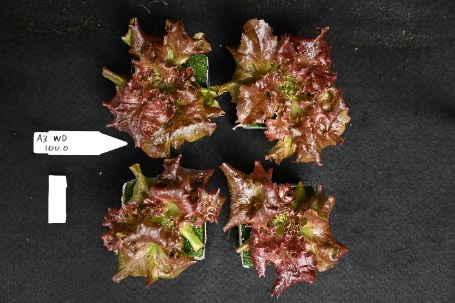

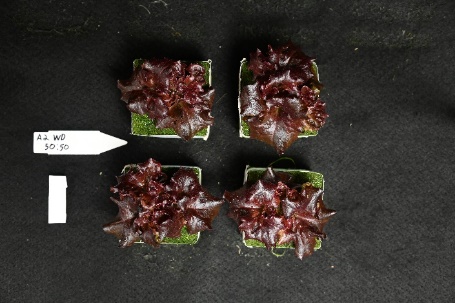

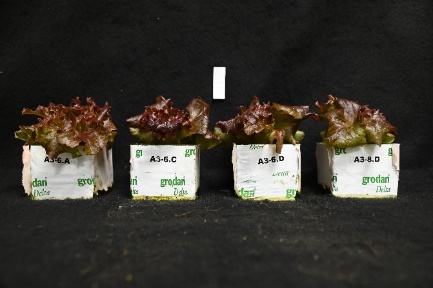

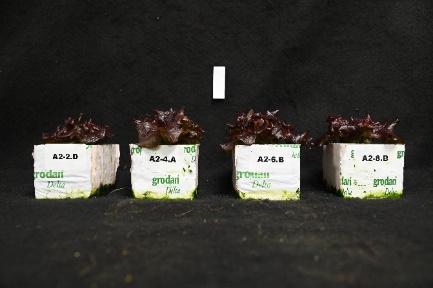

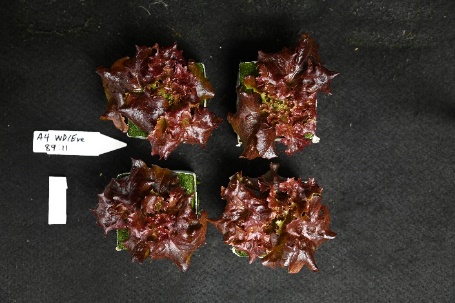

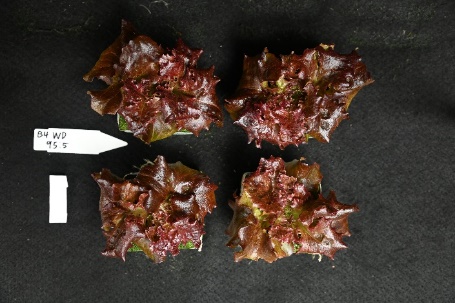

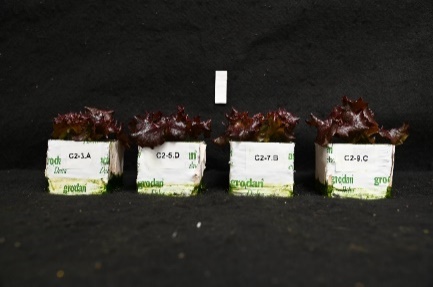

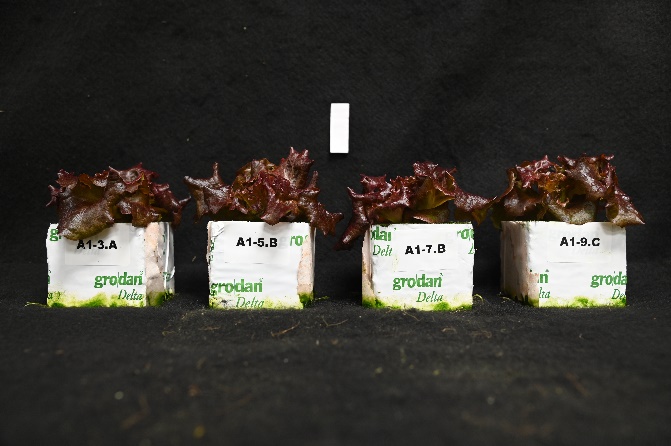

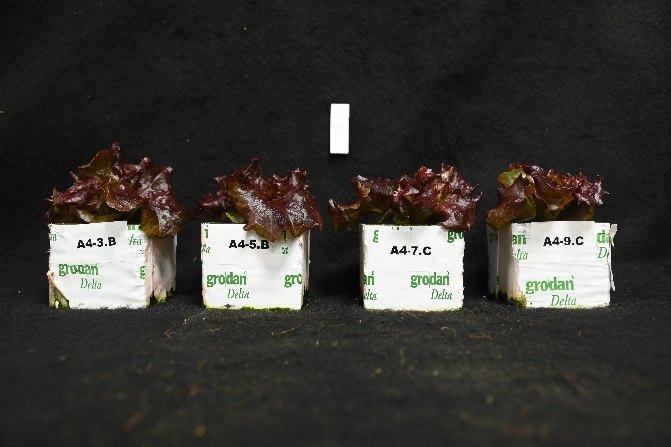

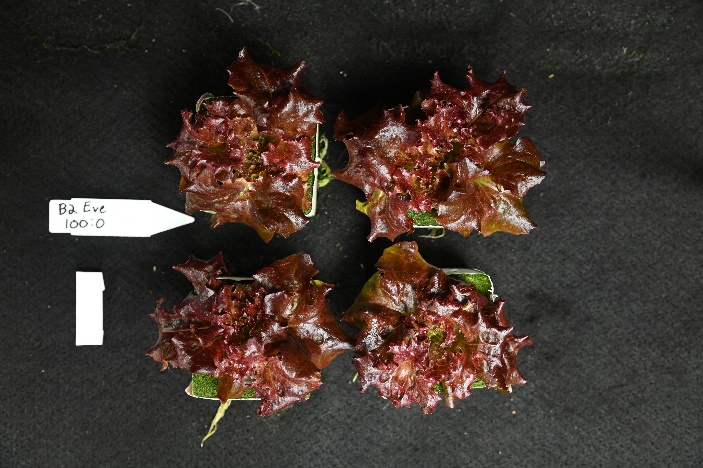

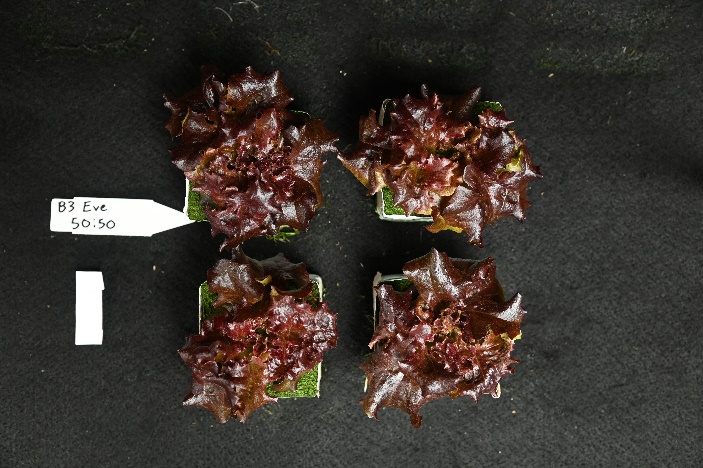

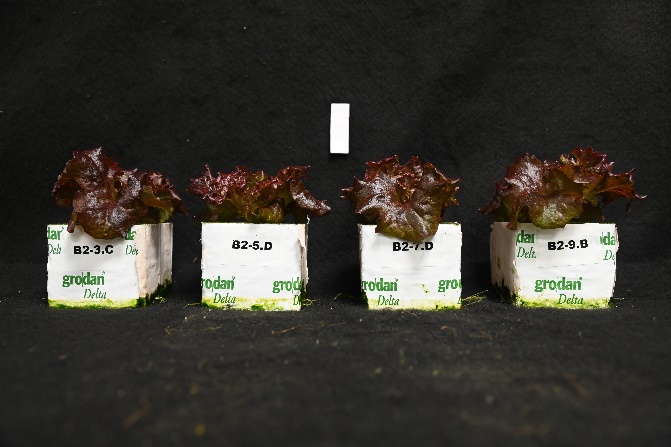

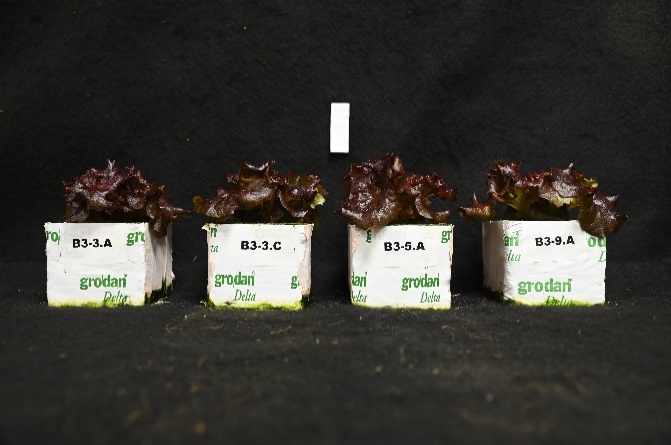

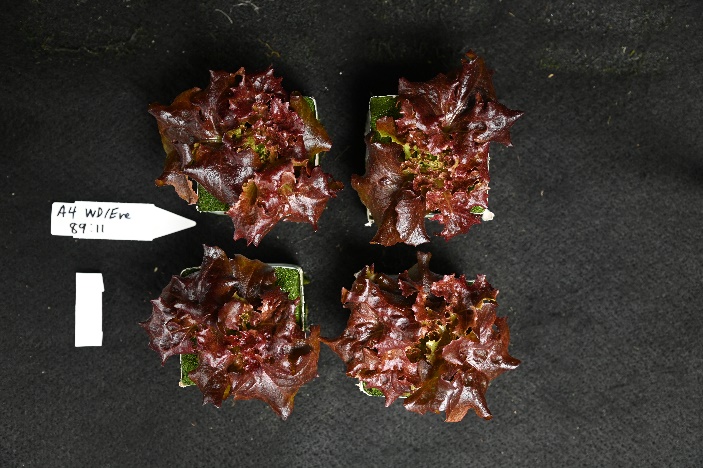

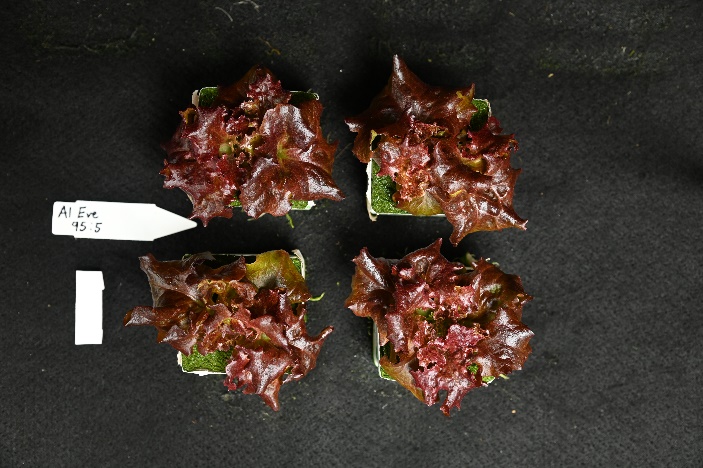

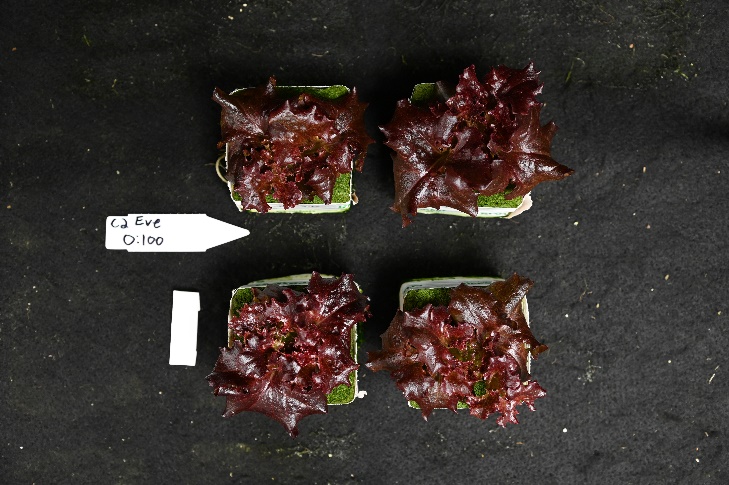

Supplemental Figure 4. Representative photos of red lettuce cv. cv. “Redflash” grown under one of five R:B*_X_* treatments for Morning, Whole Day, and Evening Treatments

Representative photos of lettuce cv. “Redflash” grown under Morning (top), Whole Day (middle), and Evening (bottom) treatments of one of five R:B_X_ treatments: (left to right) R:B_100:0_, R:B_95:5_, R:B_89:11_, R:B_50:50_, R:B_0:100_ . For all panels: upper photos are side-view representative photos, lower photos are overhead representative photos. Whole Day and Evening R:B_89:11_ plants were grown together, and hence the same treatment/photograph.

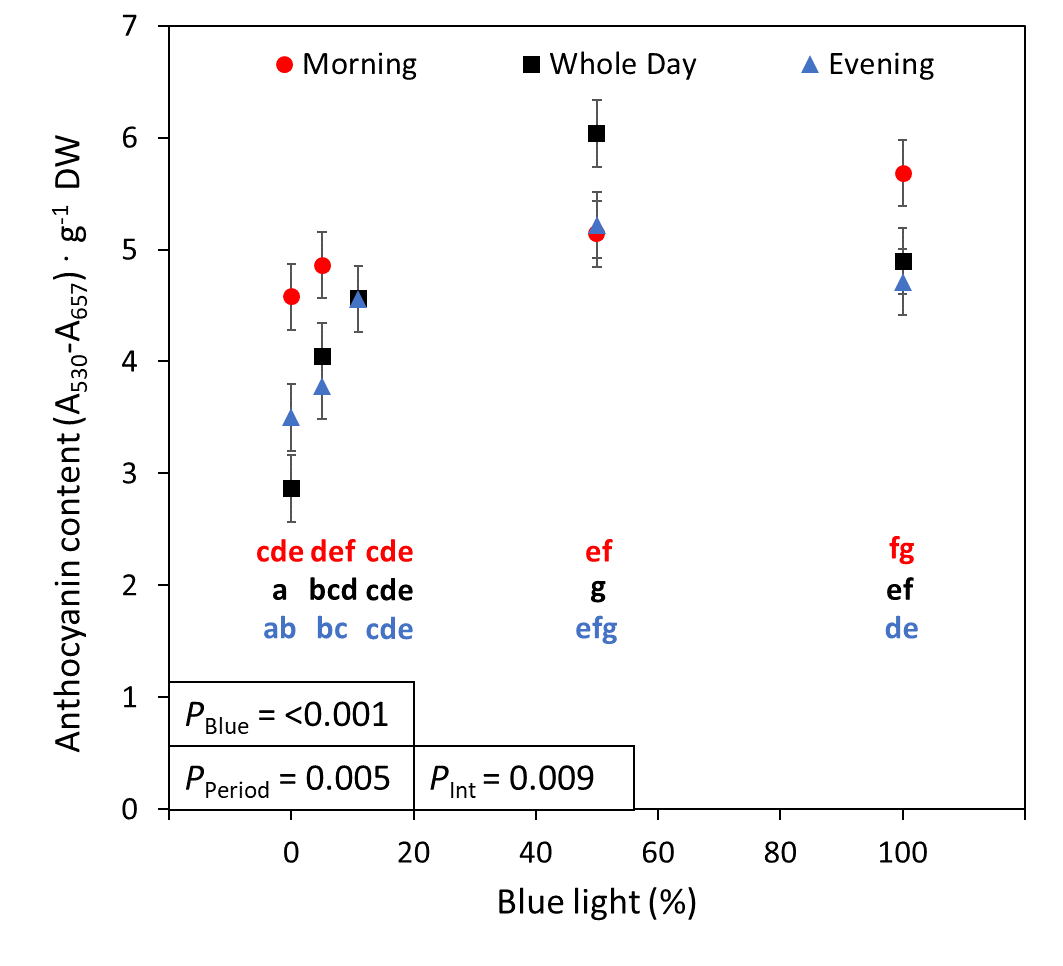

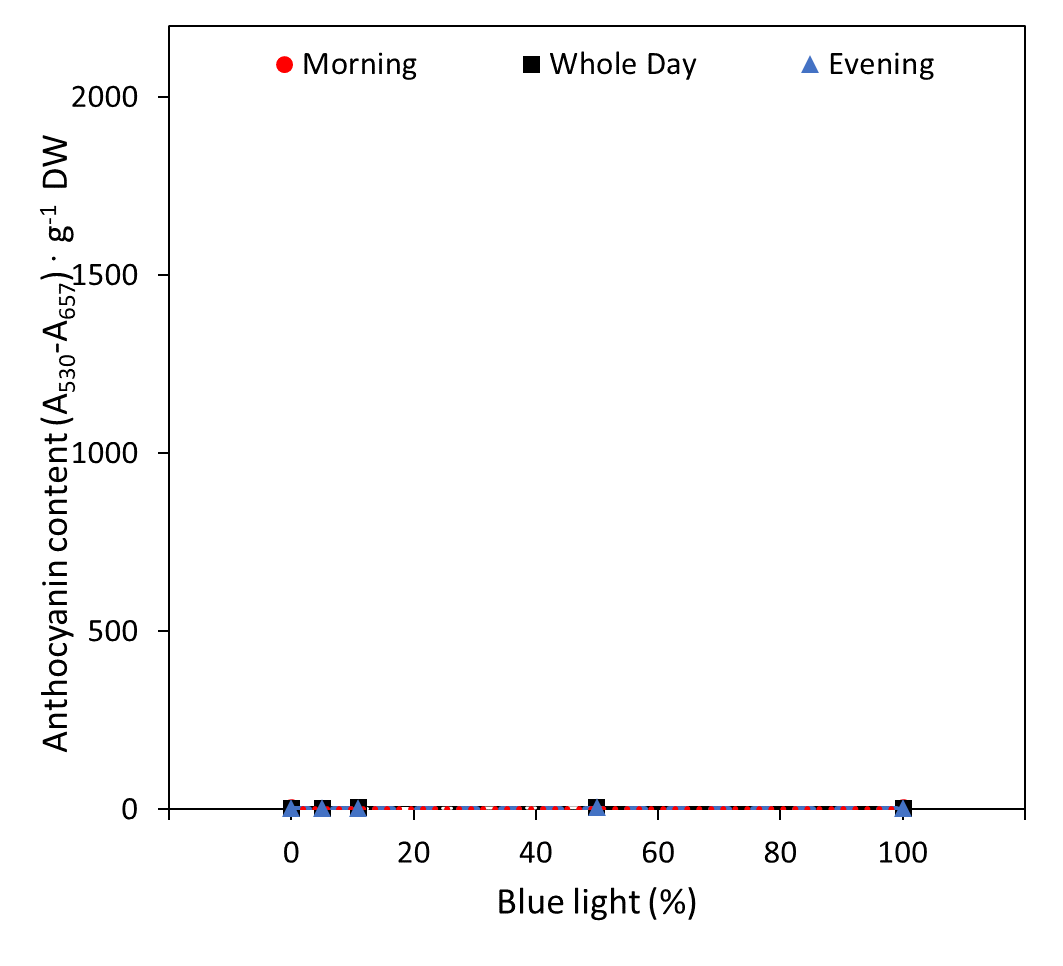

**B**

**A**

Supplemental Figure 5. Relative anthocyanin content of cv. “Greenflash” lettuce grown under five different red:blue ratios applied for three different dynamic growth periods, compared to the scale for anthocyanins in cv. “Redflash” lettuce.

For consistency, anthocyanins were measured for Greenflash lettuce, although it is a green lettuce. (**A**) shows the relative anthocyanin content of Greenflash, whereas (**B**) shows the relative anthocyanin content of Greenflash scaled to the anthocyanin content of Redflash. This is to show that Greenflash lacks anthocyanins, and the differences seen in (**A**) are likely due to trace amounts of other extracted, potentially phenolic, compounds in the material that may be present.
